# Aperiodic neural activity predicts cortical excitability in TMS-EEG

**DOI:** 10.64898/2026.10.18.759941

**Authors:** Timo van Hattem, Juliana R. Hougland, Juha Gogulski, Pedro C. Gordon, Paolo Belardinelli, Ulf Ziemann

## Abstract

**Background:** Fluctuating cortical excitability states are inherent to and critical for brain function. They can be probed non-invasively using transcranial magnetic stimulation (TMS) combined with electroencephalography (EEG) by measuring early local TMS-evoked potentials (EL-TEPs). Pre-stimulus periodic neural activity can predict fluctuating excitability states. To what extent aperiodic neural activity represents a marker of cortical excitability has not been explored.

**Objectives:** We investigated the relationship between pre-stimulus aperiodic and periodic neural activity and EL-TEP amplitude in both the primary motor cortex (M1) and dorsolateral prefrontal cortex (DLPFC).

**Methods:** Single-pulse TMS was delivered to the left M1 or left DLPFC in 29 and 11 healthy adults, respectively, while EEG was recorded simultaneously. Pre-stimulus EEG was parameterized into aperiodic and periodic components to extract the aperiodic exponent (slope of 1/f-like activity), alpha power (8–13 Hz), and beta power (13–30 Hz). EL-TEP peak-to-peak amplitudes were extracted from approximately 20-65 ms after the TMS pulse.

**Results:** Aperiodic exponent showed a strong negative relationship with EL-TEP amplitude in both M1 and DLPFC, with steeper 1/f-like slopes associated with smaller EL-TEP amplitudes. Inconsistent relationships were found between alpha and beta power with EL-TEP amplitude across the two regions.

**Conclusions:** These findings indicate that pre-stimulus aperiodic neural activity influences cortical excitability in both motor and prefrontal areas. Notably, the aperiodic exponent exhibited stronger associations with cortical excitability compared to periodic alpha or beta power. Brain-state-dependent protocols may benefit from real-time extraction of aperiodic exponent as a target feature to improve the precision and efficacy of TMS.

**Highlights:**

- Pre-stimulus aperiodic exponent predicts EL-TEP amplitude in M1 and DLPFC.
- Larger aperiodic exponent is associated with smaller EL-TEP amplitudes.
- Periodic alpha and beta power do not consistently influence EL-TEP amplitude.
- Aperiodic exponent may inform brain-state-dependent and closed-loop TMS.

## 1. Introduction

Cortical excitability refers to the reactivity of neuronal populations to internal or external stimulation. It is closely linked to the capacity of cortical circuits to undergo neuroplastic changes, which depends on the level of neuronal activity [1,2]. Transcranial magnetic stimulation (TMS) combined with electroencephalography (EEG) provides a non-invasive approach for probing cortical excitability by measuring the amplitude of TMS-evoked potentials (TEPs) [3,4]. Specifically, short-latency TEP deflections (∼15–65 ms after TMS) recorded over electrodes close to the stimulated cortex, defined as early local TEPs (EL-TEPs), have been proposed as markers of cortical excitability [5–10].

However, TMS-evoked responses exhibit substantial trial-to-trial variability [11,12]. This variability may partly reflect spontaneous fluctuations in cortical excitability over time associated with changes in the ongoing electrophysiological brain state. Indeed, a growing body of TMS-EEG research indicates that response amplitudes fluctuate as a function of the neural activity at the time of TMS [13–19]. Accounting for this by optimizing the timing of TMS delivery may enhance the precision and efficacy of TMS [20–23].

Previous TMS-EEG studies have focused on the influence of periodic (oscillatory) neural activity on TMS-evoked responses, which is characterized by narrowband frequency oscillations, such as alpha (8–13 Hz) and beta (13–30 Hz) band rhythms. Within these canonical frequency bands, phase and power are among the most commonly studied pre-stimulus EEG features for explaining fluctuations in excitability [14,18,24]. However, periodic EEG features may have limited utility as general markers of cortical excitability. Associations between oscillatory phase and power and excitability have been inconsistent, and vary substantially between and within individuals [25–27]. Moreover, reliable periodic feature extraction depends on the cortical region under investigation, potentially limiting their generalizability across the brain. Previous EEG-TMS studies have also relied predominantly on motor-evoked potentials (MEPs), which restricts the assessment of excitability to the primary motor cortex (M1). Therefore, there is a need to identify EEG features that capture and predict cortical excitability across motor and non-motor regions, as enabled by EL-TEPs.

In comparison to periodic neural activity, aperiodic neural activity has received little attention, as it has long been regarded as background neural noise. However, increasing evidence suggests that the aperiodic exponent, which reflects the slope of the broadband 1/f-like component of the EEG power spectrum, contains physiologically relevant information about the underlying state of neuronal activity [28,29]. A larger exponent (steeper slope) is believed to index lower excitability, while a smaller exponent (flatter slope) is associated with higher excitability (Figure 1) [28,30,31]. Specifically, the aperiodic exponent has recently been implicated as a marker of corticospinal excitability, demonstrating a negative relationship with MEP amplitude [32–34].

**Figure 1.**
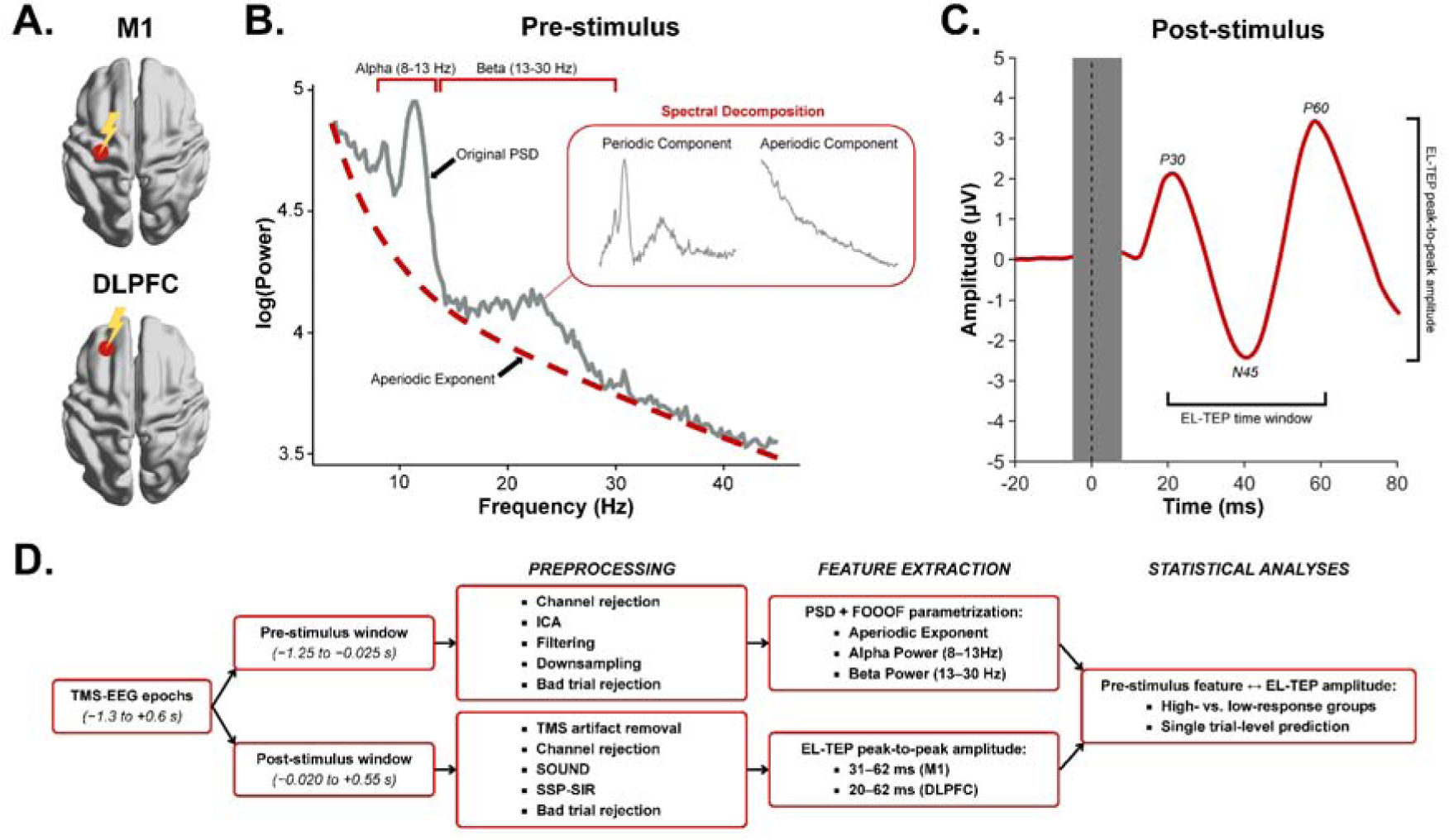
Experimental design and analytic approach. **(A)** Twenty-nine participants received 1200 single TMS pulses over left M1 while EEG was recorded simultaneously. In a separate dataset, eleven participants received 1000 single TMS pulses over the left DLPFC with concurrent EEG recording. **(B)** The power spectral density for the pre-stimulus EEG data was computed and decomposed into periodic and aperiodic components using the Fitting Oscillations & One-Over-F (FOOOF) algorithm [30]. The aperiodic exponent of the power spectrum reflects the 1/f-like slope of the aperiodic component, with a steeper slope indicating a relatively greater contribution of low-frequency activity (lower excitability) and a flatter slope indicating relatively stronger high-frequency activity (higher excitability). **(C)** Peak-to-peak amplitudes of the early local TMS-evoked potentials (EL-TEPs) were extracted from 31 to 62 ms after the pulse for M1, and 20 to 62 ms after the pulse for DLPFC (see Methods section for further details). **(D)** Analysis pipeline for preprocessing, feature extraction and statistical analyses (see Methods section for further details).

In this study, we investigated the extent to which the pre-stimulus aperiodic exponent predicts cortical excitability, indexed by EL-TEP amplitude, in both M1 and the dorsolateral prefrontal cortex (DLPFC). In addition, we examined the influence of periodic alpha and beta power on EL-TEP amplitude. We hypothesized that a larger pre-stimulus aperiodic exponent would be associated with smaller EL-TEP amplitudes, reflecting a lower cortical excitability state.

## 2. Methods

### 2.1 Participants

This study utilized two independent datasets involving TMS targeting either of the left M1 or the left DLPFC. The M1 dataset comprised 29 healthy, right-handed participants (18 female; mean age = 27.4 ± 5.1 years). The DLPFC dataset comprised 11 healthy, right-handed participants (6 female; mean age = 27.3 ± 5.1 years). Participants reported no known history of neurological or psychiatric disorders and did not have TMS or magnetic resonance imaging (MRI) contraindications. All participants provided written informed consent prior to participation. The study followed the Declaration of Helsinki and was approved by the local ethics committee of the Medical Faculty of the University of Tübingen (810/2021BO2).

### 2.2 Experimental protocols

In the M1 stimulation protocol, 1200 single TMS pulses were delivered over the hotspot of the hand representation of the left M1 at 110% of the resting motor threshold (RMT). RMT was determined as the minimum stimulation intensity, given as percentage of maximum stimulator output, that resulted in MEPs from the right first dorsal interosseous (FDI) or abductor pollicis brevis (APB) muscle with a peak-to-peak amplitude greater than 50 µV in 50% of the trials [35]. The junction of the figure-of-eight coil was oriented at approximately 45° relative to the midline. The session was divided into four blocks of 300 pulses, each block lasting approximately 15 minutes.

In the DLPFC stimulation protocol, participants received 1000 single TMS pulses over the left DLPFC (MNI coordinates: x = –32, y = 26, z = 52) [6,9,36]. The coil orientation was optimized in real-time between 45° and 90° to mitigate TMS-evoked muscle artifacts and maximize signal-to-noise in the EL-TEP time window [37]. Stimulation intensity during the experiment was set to 130% RMT, or reduced to 120% RMT if higher was not tolerable (n = 1). The session was divided into four blocks of 250 pulses, each block lasting approximately 18 minutes.

Participants were seated in a comfortable chair with their head fixed using a vacuum pillow (B.u.W Schmidt GmbH, Germany). Participants were asked to focus on a fixation cross on a monitor. Five minutes of eyes-open resting-state EEG were recorded before and after the stimulation blocks in both protocols.

### 2.3 Transcranial magnetic stimulation

Single pulses of TMS with biphasic current waveform were delivered with a MagPro R30 or X100 stimulator and figure-of-eight Cool-B65 coil (MagVenture A/S, Farum, Denmark). The randomized inter-trial interval was 2.5–3.5 s for the M1 dataset and 3.5–4.5 ms for the DLPFC dataset. Individual T1-weighted MRI images were used for neuronavigation (Localite GmbH, Germany). Individualized active noise masking was applied via earphones throughout the experiment to attenuate auditory evoked potentials from the TMS click [38].

### 2.4 EEG data acquisition

EEG was recorded using a TMS-compatible EEG cap with Ag/AgCl ring electrodes (EasyCap, Germany). Signals were sampled at 5 kHz using TMS-compatible 24-bit amplifiers (NeurOne system, Bittium Ltd., Finland). For the M1 dataset, 126 EEG channels were acquired and referenced online to FCz with ground at AFz. For the DLPFC dataset, 64 EEG channels were recorded and referenced online to the mastoid with ground at the zygomatic bone. Electrode impedances were below 5 kΩ and constantly monitored throughout the session.

### 2.5 Data preprocessing

Data preprocessing was done using MATLAB v2024b (MathWorks, Natick, MA, USA) with built-in functions from EEGLAB version 2024.2 [39] and the TESA toolbox [40,41], as well as customized scripts.

#### 2.5.1. EEG preprocessing

EEG data from concatenated blocks were epoched from –1.3 to 0.6 s around the TMS pulse and then split into pre-stimulus (–1.25 to –0.025 s) and post-stimulus (–0.020 to 0.55 s) windows. The pre- and post-stimulus windows were processed independently to preserve the causal structure of the data [13]. Detailed preprocessing steps are reported in the Supplementary Materials. A schematic overview of the preprocessing steps is shown in Figure 1.

#### 2.5.2 Pre-stimulus window: aperiodic and periodic feature extraction

The power spectral density (PSD) of the pre-stimulus EEG was computed for each trial for every electrode within the region of interest (ROI) using Welch’s method (frequency range: 4–50 Hz; window length: 1 s, 50% overlap). The M1 ROI comprised electrodes C3, C1, FC1, and FC3, whereas the DLPFC ROI included F3, F1, FC3, F5, FC1, and FC5. Aperiodic and periodic components of the mean ROI PSD were parameterized using the Fitting Oscillations & One-Over-F (FOOOF) algorithm (4–45 Hz; peak width limits: (2,6), maximum number of peaks: 4; minimum peak height: 0.1) [33,42]. Goodness-of-fit values (R²) were also retained for quality assessment. A conservative quality-control criterion was applied by excluding trials with fits where R² < 0.1 (mean ± SD rejected trials: M1 = 0.03% ± 0.05%; DLPFC = 0.07% ± 0.13%).

In addition, mean spectral power was calculated in the alpha (8–13 Hz) and beta (13–30 Hz) bands from the periodic component. Periodic alpha and beta bands were included because these frequency bands are the most commonly investigated oscillatory features in relation to cortical excitability across these two regions [16,18,43]. For DLPFC, theta power (4–8 Hz) was additionally explored given its relevance to prefrontal excitability (see Supplementary Materials for further details) [44–46].

#### 2.5.3 Post-stimulus window: peak-to-peak EL-TEP amplitude extraction

EL-TEP amplitude was quantified as the absolute peak-to-peak difference between the minimum and maximum voltage within the predefined EL-TEP time window [6,16,47]. The EL-TEP time window was defined separately for each dataset based on the grand-average TEP of all participants to account for differences in the latencies of early TEP components across stimulation sites. Peaks in the grand-average TEP were identified within predefined canonical latency windows using a prominence threshold of 0.2 μV (M1: P30 = 36 ms, N45 = 43 ms, P60 = 57 ms; DLPFC: N45 = 41 ms, P60 = 57 ms; Supplementary Figure 1) [47]. The lower and upper bounds of the EL-TEP time window were defined as 5 ms before the P30 latency and 5 ms after the P60 latency to account for inter-individual variability in peak latencies. As P30 could not be reliably identified in the DLPFC grand average, a canonical latency was used. Accordingly, the resulting EL-TEP time windows were 31–62 ms for M1 and 20–62 ms for DLPFC.

Later TEP components (> ∼100 ms) are increasingly confounded by somatosensory-evoked activity rather than reflecting cortical excitability due to TMS alone [48–50]. As a control analysis, we applied the same analytical approach used for EL-TEPs to the N100-P200 peak-to-peak amplitude (see Supplementary Materials for further details). Additionally, the M1 P60 component may be partly influenced by reafferent somatosensory feedback resulting from corticospinal activation during suprathreshold stimulation [51,52]. We therefore performed a control analysis using a restricted EL-TEP window for M1 encompassing only approximately the P30–N45 complex (EL-TEP time window = 31–48 ms; see Supplementary Materials for further details).

### 2.6 Statistical analysis

All statistical analyses were conducted in Rstudio (v4.5.1, R Foundation for Statistical Computing, Vienna, Austria), with significance set at p < 0.05. Effect sizes are reported using Cohen’s f (small = 0.10, medium = 0.25, large = 0.40) [53]. Results are presented with the mean ± standard error of the mean, unless stated otherwise.

EL-TEP amplitudes were log-transformed to improve distributional normality and reduce flooring effects. To facilitate comparison across pre-stimulus EEG features, all features were standardized using z-score normalization.

#### 2.6.1 High- versus low-response groups

To characterize the pre-stimulus brain states associated with relatively high and low cortical responses to TMS, all trials were separated within each participant according to EL-TEP amplitude and averaged within bins. For each participant, trials were split into high- and low-response groups based on the 150 highest and lowest EL-TEP amplitudes, and a grand-average waveform was generated for both groups. A subset of 150 trials per participant was selected in order to have sufficient trials to obtain reliable TEPs while retaining group-level variation in cortical excitability [36,54]. The mean pre-stimulus aperiodic exponent, alpha and beta power were calculated across the high- and low-response groups separately and compared using two-sided paired t-tests. The Benjamini Hochberg false-discovery rate (FDR) correction was used to correct for multiple comparisons [55]. Analyses were performed separately for M1 and DLPFC. The analysis was also performed on source-reconstructed EEG signals (see Supplementary Materials for further details).

#### 2.6.2 Single-trial predictions

Brain-state-dependent and closed-loop TMS paradigms generally rely on the ability to predict responses at the single-trial level. We therefore tested whether EL-TEP amplitude could also be predicted from pre-stimulus EEG features at single-trial level. A linear mixed-effects model was fit to all trials to assess the effect of pre-stimulus EEG features on the EL-TEP amplitude, with aperiodic exponent, alpha power, and beta power as fixed effects, and participant as a random intercept:

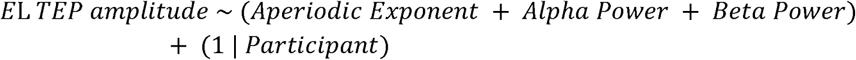

Model fits were assessed using residual diagnostics (Supplementary Figure 2). Statistical significance was validated via permutation testing (N = 5000). Predictor values were shuffled within-participants, the model refit each time, and p-values calculated as the proportion of permuted coefficients whose absolute value equaled or exceeded that of the original model [56].

Because the association between pre-stimulus brain states and EL-TEP amplitude may vary between individuals, we extended the single-trial prediction model by allowing the effects of the EEG features to vary across participants. For this, z-scored EEG features were added as random slopes in the extended model.

## 3. Results

### 3.1 Larger EL-TEP amplitudes are associated with smaller pre-stimulus aperiodic exponent

In M1, larger EL-TEP amplitudes were associated with smaller aperiodic exponent (High-response = –0.21 ± 0.05; Low-response = 0.21 ± 0.03; t(28) = –5.15, p < 0.001; Figure 2C) and lower beta power (High-response = –0.07 ± 0.03; Low-response = 0.08 ± 0.02; t(28) = – 3.02, p = 0.008; Figure 2E). No significant difference was found in pre-stimulus alpha power between the high- and low-response group (High-response = –0.04 ± 0.04; Low-response = 0.003 ± 0.03; t(28) = –0.69, p = 0.499; Figure 2D). A control analysis using a restricted EL-TEP window to the P30–N45 complex resulted in similar associations in M1 (Supplementary Figure 3).

**Figure 2.**
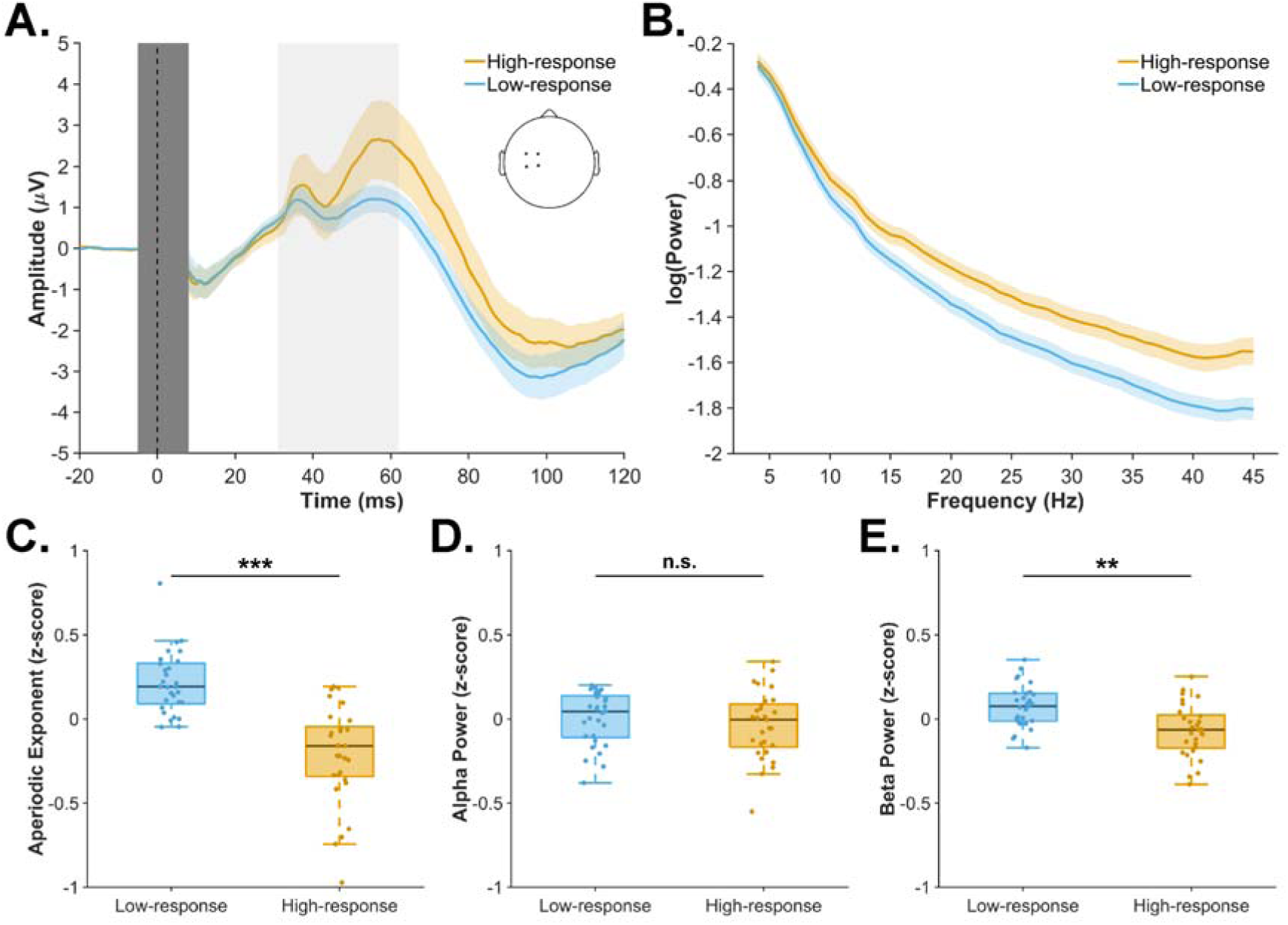
Effect of aperiodic and periodic neural activity on EL-TEP amplitude in M1. **(A)** Sensor-level grand-average waveforms across all participants (N = 29) for the high- and low-response groups, defined as the 150 highest and lowest EL-TEP trials, respectively. The ROI included electrodes C3, C1, FC1, and FC3, as indicated in the inset. The vertical light gray window represents the EL-TEP time window (31–62 ms). The vertical dark gray window represents the data affected by the TMS pulse artifact that was removed and interpolated. The coloured shaded areas around the waveform show ± SEM. **(B)** Average aperiodic fits for the high- and low-response groups. The high-response group exhibited a flatter aperiodic slope (i.e., smaller aperiodic exponent) than the low-response group. The coloured shaded areas around the aperiodic fit show ± SEM. **(C–E)** Box plots show the median, interquartile range, and whiskers extending to 1.5 times the interquartile range of the pre-stimulus EEG feature for the high- and low-response groups. Y-axes show z-score normalized pre-stimulus EEG feature values. Dots represent data from individual participants. Compared with the low-response group, the high-response group showed a significantly smaller pre-stimulus aperiodic exponent and lower beta power. Pre-stimulus alpha power was not significantly different between groups. ** p < .01; *** p < .001.

In DLPFC, larger EL-TEP amplitudes were associated with a significantly smaller pre-stimulus aperiodic exponent (High-response = –0.25 ± 0.09; Low-response = 0.22 ± 0.06; t(10) = –3.61, p = 0.014; Figure 3C). In contrast, pre-stimulus alpha power (High-response = 0.03 ± 0.08; Low-response = –0.06 ± 0.06; t(10) = 0.69, p = 0.507; Figure 3D) and beta power (High-response = 0.02 ± 0.03; Low-response = –0.04 ± 0.04; t(10) = 1.04, p = 0.484; Figure 3E) did not differ between response groups. Prefrontal theta did not influence EL-TEPs amplitude in DLPFC (Supplementary Figure 4).

**Figure 3.**
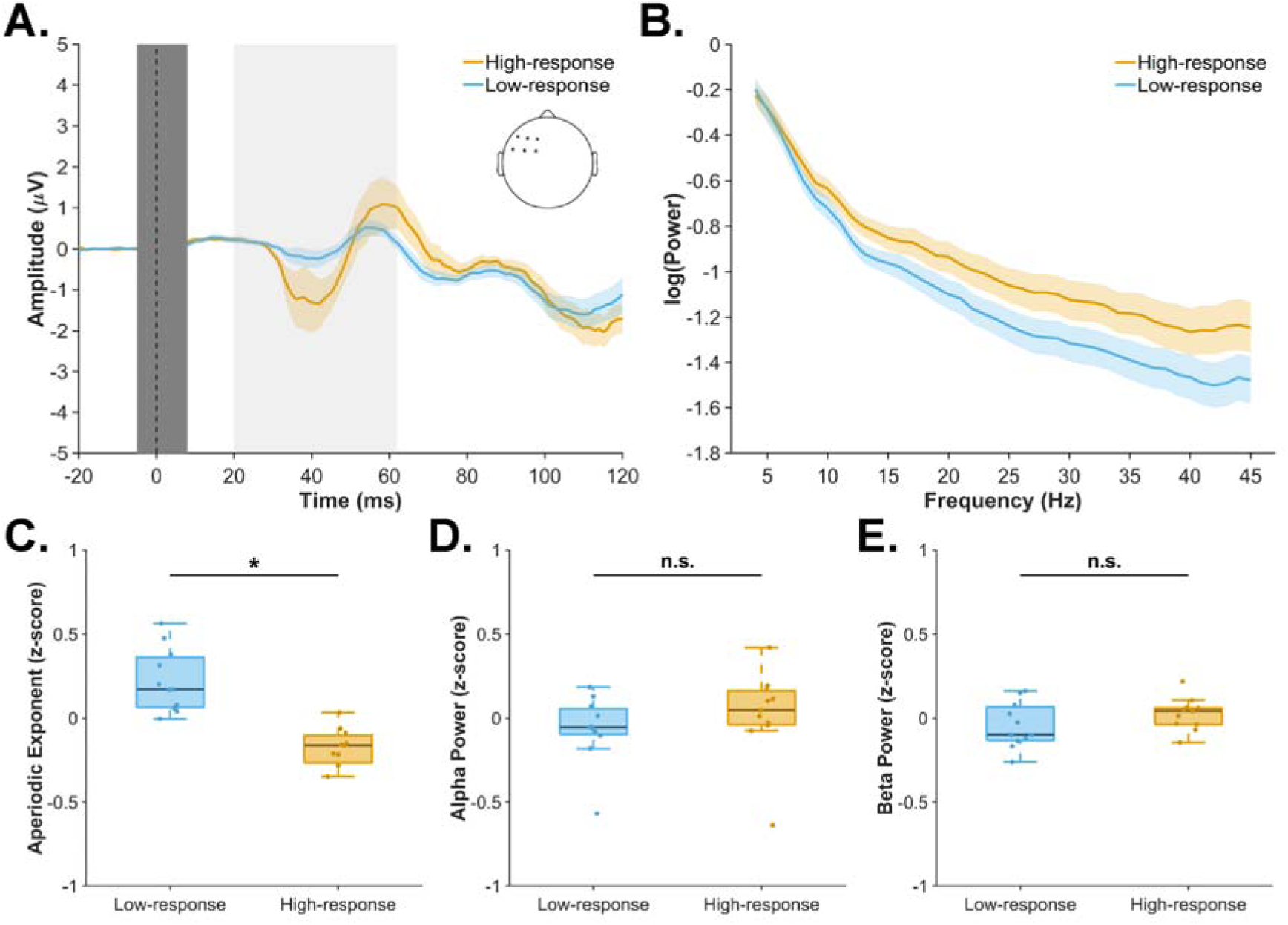
Effect of aperiodic and periodic EEG features on EL-TEP amplitude in DLPFC. **(A)** Sensor-level grand-average waveforms across all participants (N = 11) for the high- and low-response groups, defined as the 150 highest and lowest EL-TEP trials, respectively. The ROI included electrodes F3, F1, FC3, F5, FC1, and FC5, as indicated in the inset. The vertical light gray window represents the EL-TEP time window. The vertical dark gray window represents the data affected by the TMS pulse artifact that was removed and interpolated. The coloured shaded areas around the waveform show ± SEM. **(B)** Average aperiodic fits for the high- and low-response groups. The high-response group exhibited a flatter aperiodic slope (i.e., smaller aperiodic exponent) than the low-response group. The coloured shaded areas around the aperiodic fit show ± SEM. **(C–E)** Box plots show the median, interquartile range, and whiskers extending to 1.5 times the interquartile range of the pre-stimulus EEG feature for the high- and low-response groups. Y-axes show z-score normalized pre-stimulus EEG feature values. Dots represent data from individual participants. Compared with the low-response group, the high-response group showed a significantly smaller pre-stimulus aperiodic exponent. Pre-stimulus alpha and beta power did not differ significantly between groups. * p < .05.

Source-level analysis yielded a convergent pattern across both regions, with larger EL-TEP amplitudes associated with a smaller aperiodic exponent in both M1 and DLPFC (Supplementary Figure 5 and 6).

### 3.2 Aperiodic exponent predicts single-trial EL-TEP amplitude in M1 and DLPFC

In M1, aperiodic exponent (β = –0.054, SE = 0.002, t(31280) = –24.91, p < 0.001, f = 0.14; Figure 4A), alpha power (β = –0.016, SE = 0.002, t(31280) = –7.02, p < 0.001, f = 0.04; Figure 4B), and beta power (β = –0.009, SE = 0.002, t(31280) = –3.96, p < 0.001, f = 0.02; Figure 4C) were all negatively associated with EL-TEP amplitude.

**Figure 4.**
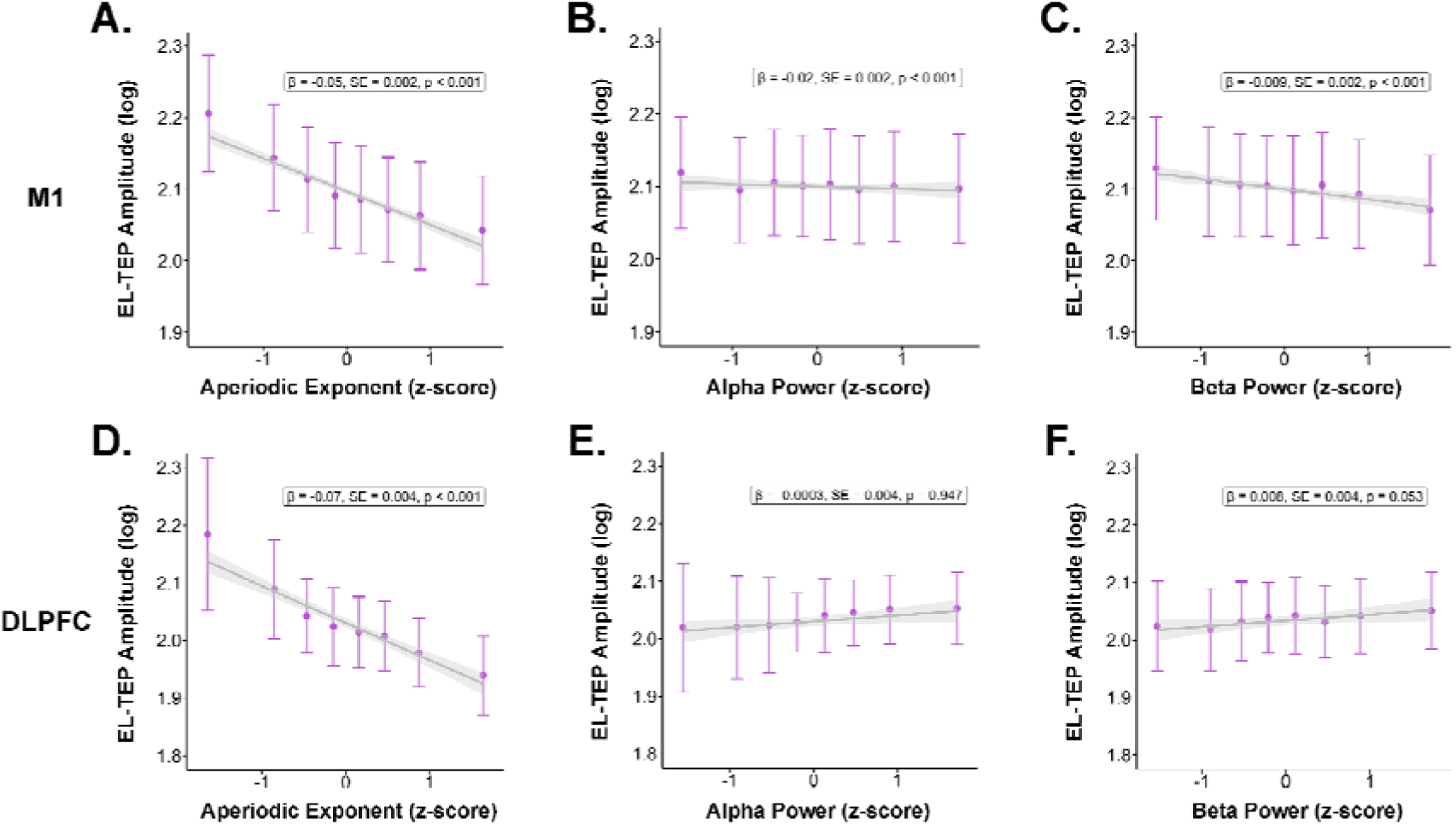
Single-trial predictions of EL-TEP amplitude using pre-stimulus EEG features in M1 and DLPFC. A linear mixed-effects model was fit to all trials to assess the effect of pre-stimulus EEG features on the EL-TEP amplitude, with aperiodic exponent, alpha power, and beta power as fixed effects, and participant as a random intercept. Grey lines represent the linear mixed-effects model-estimated relationship between each pre-stimulus EEG feature and EL-TEP amplitude. Grey shaded areas indicating 95% confidence intervals. Data points show binned mean values across octiles plotted as mean ± SEM. X-axes show z-score normalized pre-stimulus EEG feature values. Text boxes report the fixed-effect estimates (β), standard errors (SE), and p-values for each EEG feature.

In DLPFC, the linear mixed-effects model showed a significant negative association of EL-TEPs with aperiodic exponent (β = –0.074, SE = 0.004, t(9866) = –18.88, p < 0.001, f = 0.17; Figure 4D), but no significant association with alpha power (β = –0.0003, SE = 0.004, t(9866) = –0.07, p = 0.947, f < 0.01; Figure 4E) or beta power (β = 0.008, SE = 0.004, t(9866) = 1.94, p = 0.053, f = 0.02; Figure 4F).

Permutation test results supported the observed statistical significance for all EEG features in M1 (p-permutation < 0.001; Supplementary Figure 7). In DLPFC, permutation test results supported the observed statistical significance values for aperiodic exponent (p-permutation < 0.001) and alpha power (p-permutation = 0.944; Supplementary Figure 7). In contrast, beta power did narrowly reach statistical significance unlike in the original model (p-permutation = 0.049; Supplementary Figure 7).

### 3.3 Individual variability in EEG features and their associations with EL-TEP amplitude

In the extended M1 model with EEG features included as random slopes, the main effects of aperiodic exponent (β = –0.051, SE = 0.010, t(28.057) = –5.087, p < 0.001) and beta power (β = –0.010, SE = 0.003, t(28.701) = –3.021, p = 0.005) were significant, whereas alpha power was no longer significant (β = –0.011, SE = 0.007, t(28.125) = –1.515, p = 0.141). Only 62% of participants showed negative direction of the association between alpha power and EL-TEP amplitude, in comparison to 83% of participants for aperiodic exponent and 90% for beta power in M1 (Figure 5A).

**Figure 5.**
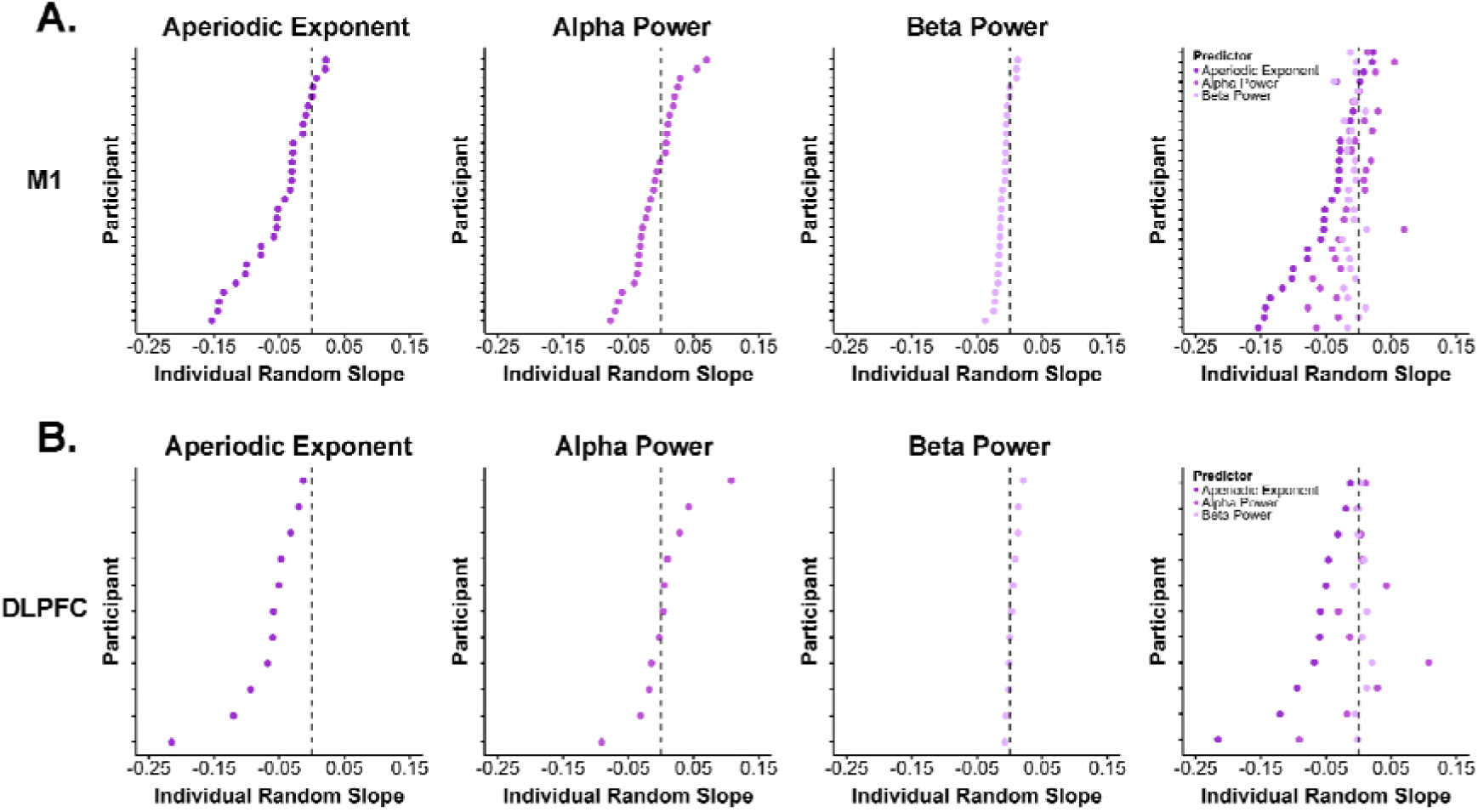
Participant-specific associations between pre-stimulus EEG and single-trial EL-TEP amplitude. The single-trial linear mixed-effects model was extended to include participant-specific random slopes for pre-stimulus EEG features, allowing the association between each EEG feature and EL-TEP amplitude to vary across participants. X-axes show the individual random slope estimates (β). The dashed line at zero indicates no association, with negative and positive values representing negative and positive associations, respectively. The Y-axes denote participants, with each tick representing one participant. Each dot represents the participant-specific random slope for a given pre-stimulus EEG feature. In the first three columns, participant-specific slopes are ranked from highest to lowest separately for each EEG feature so the vertical position of a participant does not correspond across these columns. In the final column, participants are aligned across EEG features, such that each row represents the same participant and shows their corresponding slopes for aperiodic exponent, alpha power, and beta power.

In the DLPFC, the effects were significant for aperiodic exponent (β = –0.071, SE = 0.018, t(10.087) = –3.921, p = 0.003), but not alpha power (β = 0.004, SE = 0.016, t(9.998) = 0.246, p = 0.811), and beta power (β = 0.004, SE = 0.005, t(9.397) = 0.743, p = 0.476). All participants (100%) showed negative slopes for aperiodic exponent, whereas only 45% showed positive slopes for both alpha and beta power (Figure 5B).

Across both datasets, between-participant variability in aperiodic exponent (CV M1 = 21.3%; CV DLPFC = 36.8%) was lower than within-participant variability (CV M1 = 37.1%; CV DLPFC = 49.2%). Similar to the aperiodic exponent, within-participant variability exceeded between-participant variability for both alpha (within-participant CV M1 = 59.5%, between-participant CV M1 = 40.7%; within-participant CV DLPFC = 72.1%, between-participant CV DLPFC = 37.2%) and beta power (within-participant CV M1 = 44.7%, between-participant CV M1 = 21.7%; within-participant CV DLPFC = 48.9%, between-participant CV DLPFC = 33.5%).

## 4. Discussion

The present study investigated whether cortical excitability in M1 and DLPFC is modulated by pre-stimulus aperiodic and periodic neural activity. We found that states of lower cortical excitability, indexed by smaller EL-TEP amplitude, were associated with larger pre-stimulus aperiodic exponents in both cortical areas. In contrast, the relationship between EL-TEP amplitude with periodic alpha and beta power was less consistent and region-dependent. Taken together, our findings suggest that aperiodic neural activity reflects cortical excitability states across distinct cortical regions.

### 4.1 Aperiodic exponent reflects cortical excitability states

A number of recent studies have demonstrated that lower corticospinal excitability, as reflected by smaller peak-to-peak MEP amplitude, was associated with larger pre-stimulus aperiodic exponent [32–34]. In contrast to MEPs, TEPs are considered direct measures of cortical excitability and can be assessed also in non-motor regions [57]. Our findings on EL-TEPs aligned with those on MEPs: smaller peak-to-peak amplitude was associated with larger pre-stimulus aperiodic exponent. Importantly, the relationship between aperiodic exponent and EL-TEP amplitude was observed both in M1 and DLPFC, suggesting that the relationship may generalize beyond the motor system.

Although the physiological interpretation of the aperiodic exponent remains under active investigation, accumulating evidence suggests that it reflects aspects of neuronal excitability. Previous research has suggested that the aperiodic exponent measures excitation/inhibition (E/I) balance, which reflects the ratio of faster glutamate (excitatory) and slower GABAergic (inhibitory) synaptic currents, where a steeper slope means more inhibition and a flatter slope means more excitation [31]. At the level of the EL-TEP, our findings are broadly consistent with the E/I balance hypothesis, in which larger pre-stimulus aperiodic exponents were related to smaller EL-TEPs. Simply, a higher inhibitory state elicited lower cortical excitability responses.

Although E/I balance is a common interpretation of the aperiodic exponent, there are some limitations to this hypothesis [28,29]. For example, pharmacological manipulation with GABA-A receptor antagonists did not modulate aperiodic exponent in mice [58]. In addition, the phase of the sensorimotor mu-alpha rhythm interacts with the effect of aperiodic exponent on MEP amplitudes, indicating that the aperiodic exponent may not represent a strict one-to-one mapping of E/I balance [33]. Rather, the aperiodic exponent likely reflects multiple underlying physiological mechanisms, which may include E/I balance to an extent [28,29]. Another potential mechanism is the communication-through-aperiodic-dynamics hypothesis which proposes that steeper 1/f-like activity reflects a shift towards slower dominant timescales which are associated with facilitated network communication [29,59]. While our findings help to contextualize the influence of aperiodic neural activity on TMS-evoked responses, further research is necessary to elucidate the underlying physiological mechanisms.

It is important to note that larger EL-TEP amplitude may not necessarily be a marker of higher cortical *excitation*. Rather, larger EL-TEP amplitudes are representative of cortical responsiveness (*excitability*), which could be mediated by both excitatory and inhibitory mechanisms [60–62]. In this study, we looked at the EL-TEP as a general readout of excitability which encompasses a relatively broad latency window. As the EL-TEP includes TEP deflections from ∼20 to 60 ms (P30, N45, P60), neurophysiological interpretations with regard to engagement of distinct excitatory and inhibitory populations across different cortical regions based on this outcome is limited.

Notably, pre-stimulus EEG features did not predict the amplitude of later TEP components, which are believed to reflect primarily reafferent somatosensory processing (Supplementary Figures 3, 8, and 9) [48,50,52]. This lack of association suggests that the observed relationships are more specific to early TMS-evoked cortical responses, supporting their interpretation as markers of cortical excitability.

### 4.2 Modulation of cortical excitability by periodic neural activity

The high-response group in M1 was characterized by lower pre-stimulus beta power compared to the low-response group, while there was no significant difference in pre-stimulus alpha power between groups. At the level of single-trial responses, we observed a negative association between both alpha and beta power and EL-TEP amplitude in the original model. However, the association between EL-TEP amplitude and alpha power was no longer significant after allowing the relationship to vary across participants, indicating substantial inter-individual heterogeneity in this association. Although the exact role of these rhythms in relation to excitability is not fully understood, sensorimotor alpha and beta activity have been linked to gating GABAergic inhibitory processes [14,63–66].

In DLPFC, pre-stimulus alpha and beta power did not significantly differ between high- and low-response groups. At the single-trial level, we observed only a weak positive effect of beta power on EL-TEP amplitude in the original model, but the extended model suggests that this was also driven by substantial inter-individual heterogeneity. Together, these findings suggest that the relationship between periodic neural activity and cortical excitability may not be consistent across participants and not uniform across cortical regions. This variability may reflect differences in the functional and neurophysiological relevance of alpha and beta oscillations depending on the stimulated target and the underlying cortical circuitry.

In contrast to the current findings, previous work has reported positive associations between periodic alpha and beta power and TEP amplitude across M1 and DLPFC [13,16,67]. Methodological differences between studies investigating the effects of brain states on TEP amplitude may partly account for conflicting findings, including the TEP quantification approach, EEG preprocessing procedure, periodic and aperiodic spectral component separation, and statistical analysis strategy [43]. Single-trial TEP amplitudes have a relatively low signal-to-noise and are highly susceptible to data quality and processing parameters, which may contribute to inconsistencies across studies.

### 4.3 Implications for brain-state-dependent neuromodulation

An emerging field of EEG-TMS research focuses on the development of brain-state-dependent and closed-loop TMS protocols, which could help control the variability in responses to TMS, thereby potentially increasing the consistency of stimulation effects across trials and sessions [22,23,68–70]. Delivering stimulation specifically during high or low excitability states has been shown to increase the efficacy of TMS-induced plasticity [19,45,71,72].

In previous brain-state-dependent TMS studies, stimulation timing has been exclusively determined by periodic EEG features, such as power and phase of the sensorimotor mu-rhythm or midfrontal theta [19,45]. However, the reported effects of periodic power and phase on excitability are inconsistent and variable both between and within participants [26,27,33,73]. Our findings indicate that the association between aperiodic exponent and EL-TEP amplitude is more consistent across individuals and stimulation targets than alpha and beta power, which could improve practical implementation and reduce effect heterogeneity in TMS effects. The effect size of the single-trial relationship for aperiodic exponent with EL-TEP amplitude was approximately 300% to 900% larger than for alpha or beta power in both M1 and DLPFC, suggesting that the aperiodic exponent may be a stronger predictor of cortical excitability.

### 4.4 Limitations

Because TEPs are embedded within ongoing EEG activity, a potential concern is that differences in EL-TEP amplitude may in part reflect differences in ongoing pre-stimulus neural dynamics that persist into the post-stimulus period [13,15,16,67]. No established approach currently allows these contributions to be fully disentangled. We performed a control analysis to assess whether comparable associations could be observed in the absence of TMS-evoked activity by inserting artificial triggers in resting-state EEG (see Supplementary Materials and Supplementary Figure 10 for further details). Although we found no significant associations between the pre-stimulus aperiodic exponent or periodic alpha and beta power and the corresponding post-trigger EEG amplitudes, we cannot completely exclude interactions between ongoing activity and the TMS-evoked response.

The aperiodic and periodic EEG components arise from activity in the same underlying neural activity and are parametrized from the same power spectrum. Thus, aperiodic exponent and periodic power cannot be fully dissociated and interpretation as completely independent features warrants caution.

Finally, the DLPFC analysis was based on a relatively small sample size. Replication with larger sample size will be important to determine the reliability and generalizability of the DLPFC findings.

## 5. Conclusions

This study demonstrates that aperiodic neural activity provides an index of cortical excitability across motor and non-motor cortical regions. Compared with periodic EEG features, such as alpha and beta power, the aperiodic exponent showed stronger and more consistent associations with EL-TEP amplitude, suggesting that it captures information about the excitability state that is not fully explained by oscillatory activity. Together, these findings strongly support the potential of the aperiodic exponent as a target feature for brain-state-dependent neuromodulation across cortical regions, which could enhance the precision and efficacy of TMS interventions.

## Data availability

Data is available upon reasonable request and after signing a data sharing agreement.

## CRediT authorship contribution statement

**Timo van Hattem**: Conceptualization, Methodology, Investigation, Data Curation, Formal analysis, Visualization, Writing – original draft, Writing – review & editing. **Juliana R. Hougland**: Conceptualization, Methodology, Formal analysis, Visualization, Writing – original draft, Writing – review & editing. **Juha Gogulski**: Writing – review & editing. **Pedro C. Gordon**: Writing – review & editing. **Paolo Belardinelli**: Formal analysis, Writing – review & editing. **Ulf Ziemann**: Supervision, Funding acquisition, Resources, Writing – review & editing.

## References

1. Bienenstock, E., Cooper, L., & Munro, P. (1982). Theory for the development of neuron selectivity: Orientation specificity and binocular interaction in visual cortex. The Journal of Neuroscience, 2(1), 32–48. 10.1523/JNEUROSCI.02-01-00032.1982

2. Huang, Y.-Z., Lu, M.-K., Antal, A., Classen, J., Nitsche, M., Ziemann, U., Ridding, M., Hamada, M., Ugawa, Y., Jaberzadeh, S., Suppa, A., Paulus, W., & Rothwell, J. (2017). Plasticity induced by non-invasive transcranial brain stimulation: A position paper. Clinical Neurophysiology, 128(11), 2318–2329. 10.1016/j.clinph.2017.09.007

3. Hernandez-Pavon, J. C., Veniero, D., Bergmann, T. O., Belardinelli, P., Bortoletto, M., Casarotto, S., Casula, E. P., Farzan, F., Fecchio, M., Julkunen, P., Kallioniemi, E., Lioumis, P., Metsomaa, J., Miniussi, C., Mutanen, T. P., Rocchi, L., Rogasch, N. C., Shafi, M. M., Siebner, H. R., … Ilmoniemi, R. J. (2023). TMS combined with EEG: Recommendations and open issues for data collection and analysis. Brain Stimulation, 16(2), 567–593. 10.1016/j.brs.2023.02.009

4. Ziemann, U., Bai, Y., Baumer, F. M., Beck, M. M., Belardinelli, P., Belvisi, D., Bender, S., Bergmann, T. O., Bortoletto, M., Casarotto, S., Casula, E., Chaves, A. R., De Andrade, D. C., Conte, A., Daskalakis, Z. J., Farzan, F., Ferrarelli, F., Fitzgerald, P. B., Gordon, P. C., … Ilmoniemi, R. J. (2026). Clinical utility and prospective of TMS– EEG: Updated review from an international expert group. Clinical Neurophysiology, 184, 2111487. 10.1016/j.clinph.2025.2111487

5. Chen, N.-F., Hassan, U., Ross, J. M., Forman, L., Hartford, J. W., Gogulski, J., Parmigiani, S., Truong, J., Keller, C. J., & Cline, C. C. (2025). Noninvasive profiling of input-output excitability curves in human prefrontal cortex. Neuroscience. 10.1101/2025.09.26.678876

6. Gogulski, J., Cline, C. C., Ross, J. M., Truong, J., Sarkar, M., Parmigiani, S., & Keller, C. J. (2024). Mapping cortical excitability in the human dorsolateral prefrontal cortex. Clinical Neurophysiology, 164, 138–148. 10.1016/j.clinph.2024.05.008

7. Hassan, U., Forman, L., Hartford, J. W., Lee, H., Parmigiani, S., Chen, N.-F., Ross, J. M., Solomon, E. A., Gogulski, J., Kim, N., Truong, J., Cline, C. C., & Keller, C. J. (2026). Cognitive load modulates cortical excitability in the human prefrontal cortex. Neuroscience. 10.64898/2026.04.25.720856

8. Keller, C. J., Huang, Y., Herrero, J. L., Fini, M. E., Du, V., Lado, F. A., Honey, C. J., & Mehta, A. D. (2018). Induction and Quantification of Excitability Changes in Human Cortical Networks. The Journal of Neuroscience, 38(23), 5384–5398. 10.1523/JNEUROSCI.1088-17.2018

9. Parmigiani, S., Cline, C. C., Sarkar, M., Forman, L., Truong, J., Ross, J. M., Gogulski, J., & Keller, C. J. (2025). Real-time optimization to enhance noninvasive cortical excitability assessment in the human dorsolateral prefrontal cortex. Clinical Neurophysiology, 174, 225–234. 10.1016/j.clinph.2025.02.261

10. Ross, J. M., Cline, C. C., Sarkar, M., Truong, J., & Keller, C. J. (2023). Neural effects of TMS trains on the human prefrontal cortex. Scientific Reports, 13(1), 22700. 10.1038/s41598-023-49250-7

11. Guerra, A., López-Alonso, V., Cheeran, B., & Suppa, A. (2020). Variability in non-invasive brain stimulation studies: Reasons and results. Neuroscience Letters, 719, 133330. 10.1016/j.neulet.2017.12.058

12. Ziemann, U., & Siebner, H. R. (2015). Inter-subject and Inter-session Variability of Plasticity Induction by Non-invasive Brain Stimulation: Boon or Bane? Brain Stimulation, 8(3), 662–663. 10.1016/j.brs.2015.01.409

13. Ahola, O., Haxel, L., Ermolova, M., Humaidan, D., Mutanen, T. P., Laine, M., Makkonen, M., Ukharova, E., Roine, T., Lioumis, P., Guidotti, R., Llmoniemi, R. J., & Ziemann, U. (2025). Predictive modeling of TMS-evoked responses: Unraveling instantaneous excitability states. NeuroImage, 322, 121553. 10.1016/j.neuroimage.2025.121553

14. Bergmann, T. O., Lieb, A., Zrenner, C., & Ziemann, U. (2019). Pulsed Facilitation of Corticospinal Excitability by the Sensorimotor μ-Alpha Rhythm. The Journal of Neuroscience, 39(50), 10034–10043. 10.1523/JNEUROSCI.1730-19.2019

15. Desideri, D., Zrenner, C., Ziemann, U., & Belardinelli, P. (2019). Phase of sensorimotor μ oscillation modulates cortical responses to transcranial magnetic stimulation of the human motor cortex. The Journal of Physiology, 597(23), 5671– 5686. 10.1113/JP278638

16. Perera, N. D., Wischnewski, M., Alekseichuk, I., Shirinpour, S., & Opitz, A. (2024). State-Dependent Motor Cortex Stimulation Reveals Distinct Mechanisms for Corticospinal Excitability and Cortical Responses. eNeuro, 11(11), ENEURO.0450-24.2024. 10.1523/ENEURO.0450-24.2024

17. Schaworonkow, N., Triesch, J., Ziemann, U., & Zrenner, C. (2019). EEG-triggered TMS reveals stronger brain state-dependent modulation of motor evoked potentials at weaker stimulation intensities. Brain Stimulation, 12(1), 110–118. 10.1016/j.brs.2018.09.009

18. Wischnewski, M., Haigh, Z. J., Shirinpour, S., Alekseichuk, I., & Opitz, A. (2022). The phase of sensorimotor mu and beta oscillations has the opposite effect on corticospinal excitability. Brain Stimulation, 15(5), 1093–1100. 10.1016/j.brs.2022.08.005

19. Zrenner, C., Desideri, D., Belardinelli, P., & Ziemann, U. (2018). Real-time EEG-defined excitability states determine efficacy of TMS-induced plasticity in human motor cortex. Brain Stimulation, 11(2), 374–389. 10.1016/j.brs.2017.11.016

20. Humaidan, D., Xu, J., Kirchhoff, M., Romani, G. L., Ilmoniemi, R. J., & Ziemann, U. (2024). Towards real-time EEG–TMS modulation of brain state in a closed-loop approach. Clinical Neurophysiology, 158, 212–217. 10.1016/j.clinph.2023.12.006

21. Miniussi, C., & Bortoletto, M. (2025). Harnessing neural variability: Implications for brain research and non-invasive brain stimulation. Neuroscience & Biobehavioral Reviews, 176, 106312. 10.1016/j.neubiorev.2025.106312

22. Rabuffo, G., Angiolelli, M., Fukai, T., Deco, G., Sorrentino, P., & Momi, D. (2026). Pre-stimulus brain states predict and control variability in stimulation responses. Brain Stimulation, 19(4), 103118. 10.1016/j.brs.2026.103118

23. Zrenner, C., & Ziemann, U. (2024). Closed-Loop Brain Stimulation. Biological Psychiatry, 95(6), 545–552. 10.1016/j.biopsych.2023.09.014

24. Hussain, S. J., Claudino, L., Bönstrup, M., Norato, G., Cruciani, G., Thompson, R., Zrenner, C., Ziemann, U., Buch, E., & Cohen, L. G. (2019). Sensorimotor Oscillatory Phase–Power Interaction Gates Resting Human Corticospinal Output. Cerebral Cortex, 29(9), 3766–3777. 10.1093/cercor/bhy255

25. Hougland, J. R., Kirchhoff, M., Vetter, D. E., Ahola, O., Jooß, A., Humaidan, D., & Ziemann, U. (2025). Fluctuations in the optimal sensorimotor mu-rhythm phase associated with high corticospinal excitability during TMS-EEG. Brain Stimulation, 18(6), 1843–1851. 10.1016/j.brs.2025.09.019

26. Karabanov, A. N., Madsen, K. H., Krohne, L. G., & Siebner, H. R. (2021). Does pericentral mu-rhythm “power” corticomotor excitability? – A matter of EEG perspective. Brain Stimulation, 14(3), 713–722. 10.1016/j.brs.2021.03.017

27. Madsen, K. H., Karabanov, A. N., Krohne, L. G., Safeldt, M. G., Tomasevic, L., & Siebner, H. R. (2019). No trace of phase: Corticomotor excitability is not tuned by phase of pericentral mu-rhythm. Brain Stimulation, 12(5), 1261–1270. 10.1016/j.brs.2019.05.005

28. Donoghue, T. (2025). A Systematic Review of Aperiodic Neural Activity in Clinical Investigations. European Journal of Neuroscience, 62(7), e70255. 10.1111/ejn.70255

29. Preston, M., Smith, S., & Voytek, B. (2026). Potential mechanisms and functional significance of aperiodic neural activity. Nature Human Behaviour, 10(8), 1431–1440. 10.1038/s41562-026-02503-7

30. Donoghue, T., Haller, M., Peterson, E. J., Varma, P., Sebastian, P., Gao, R., Noto, T., Lara, A. H., Wallis, J. D., Knight, R. T., Shestyuk, A., & Voytek, B. (2020). Parameterizing neural power spectra into periodic and aperiodic components. Nature Neuroscience, 23(12), 1655–1665. 10.1038/s41593-020-00744-x

31. Gao, R., Peterson, E. J., & Voytek, B. (2017). Inferring synaptic excitation/inhibition balance from field potentials. NeuroImage, 158, 70–78. 10.1016/j.neuroimage.2017.06.078

32. Frohlich, J., Ruch, S., Trunk, B. H., Keute, M., Mediano, P. A. M., & Gharabaghi, A. (2024). Brain signal complexity and aperiodicity predict human corticospinal excitability. Neuroscience. 10.1101/2024.02.09.579457

33. Hougland, J. R., Kirchhoff, M., Van Hattem, T., Rösch, J., Chen, J., Schaier, M., Belardinelli, P., & Ziemann, U. (2026). Aperiodic 1/f brain activity is associated with corticospinal excitability. NeuroImage, 339, 122162. 10.1016/j.neuroimage.2026.122162

34. Jin, J., Wang, X., Zhao, X., Wang, H., Li, Y., Liu, Z., & Yin, T. (2026). Aperiodic Brain Activity Modulates Corticospinal Excitability. Neuroscience Bulletin, 42(7), 1507– 1520. 10.1007/s12264-025-01559-1

35. Rossini, P. M., Burke, D., Chen, R., Cohen, L. G., Daskalakis, Z., Di Iorio, R., Di Lazzaro, V., Ferreri, F., Fitzgerald, P. B., George, M. S., Hallett, M., Lefaucheur, J. P., Langguth, B., Matsumoto, H., Miniussi, C., Nitsche, M. A., Pascual-Leone, A., Paulus, W., Rossi, S., … Ziemann, U. (2015). Non-invasive electrical and magnetic stimulation of the brain, spinal cord, roots and peripheral nerves: Basic principles and procedures for routine clinical and research application. An updated report from an I.F.C.N. Committee. Clinical Neurophysiology, 126(6), 1071–1107. 10.1016/j.clinph.2015.02.001

36. Gogulski, J., Cline, C. C., Ross, J. M., Parmigiani, S., & Keller, C. J. (2024). Reliability of the TMS-evoked potential in dorsolateral prefrontal cortex. Cerebral Cortex, 34(4), bhae130. 10.1093/cercor/bhae130

37. Casarotto, S., Fecchio, M., Rosanova, M., Varone, G., D’Ambrosio, S., Sarasso, S., Pigorini, A., Russo, S., Comanducci, A., Ilmoniemi, R. J., & Massimini, M. (2022). The rt-TEP tool: Real-time visualization of TMS-Evoked Potentials to maximize cortical activation and minimize artifacts. Journal of Neuroscience Methods, 370, 109486. 10.1016/j.jneumeth.2022.109486

38. Russo, S., Sarasso, S., Puglisi, G. E., Dal Palù, D., Pigorini, A., Casarotto, S., D’Ambrosio, S., Astolfi, A., Massimini, M., Rosanova, M., & Fecchio, M. (2022). TAAC - TMS Adaptable Auditory Control: A universal tool to mask TMS clicks. Journal of Neuroscience Methods, 370, 109491. 10.1016/j.jneumeth.2022.109491

39. Delorme, A., & Makeig, S. (2004). EEGLAB: An open source toolbox for analysis of single-trial EEG dynamics including independent component analysis. Journal of Neuroscience Methods, 134(1), 9–21. 10.1016/j.jneumeth.2003.10.009

40. Mutanen, T. P., Biabani, M., Sarvas, J., Ilmoniemi, R. J., & Rogasch, N. C. (2020). Source-based artifact-rejection techniques available in TESA, an open-source TMS– EEG toolbox. Brain Stimulation, 13(5), 1349–1351. 10.1016/j.brs.2020.06.079

41. Rogasch, N. C., Sullivan, C., Thomson, R. H., Rose, N. S., Bailey, N. W., Fitzgerald, P. B., Farzan, F., & Hernandez-Pavon, J. C. (2017). Analysing concurrent transcranial magnetic stimulation and electroencephalographic data: A review and introduction to the open-source TESA software. NeuroImage, 147, 934–951. 10.1016/j.neuroimage.2016.10.031

42. Suresh, T., & Hussain, S. J. (2023). Re-evaluating the contribution of sensorimotor mu rhythm phase and power to human corticospinal output: A replication study. Brain Stimulation, 16(3), 936–938. 10.1016/j.brs.2023.05.022

43. Bigoni, C., Pagnamenta, S., Cadic-Melchior, A., Bevilacqua, M., Harquel, S., Raffin, E., & Hummel, F. C. (2024). MEP and TEP features variability: Is it just the brain-state? Journal of Neural Engineering, 21(1), 016011. 10.1088/1741-2552/ad1dc2

44. Gordon, P. C., Dörre, S., Belardinelli, P., Stenroos, M., Zrenner, B., Ziemann, U., & Zrenner, C. (2021). Prefrontal Theta-Phase Synchronized Brain Stimulation With Real-Time EEG-Triggered TMS. Frontiers in Human Neuroscience, 15, 691821. 10.3389/fnhum.2021.691821

45. Gordon, P. C., Belardinelli, P., Stenroos, M., Ziemann, U., & Zrenner, C. (2022). Prefrontal theta phase-dependent rTMS-induced plasticity of cortical and behavioral responses in human cortex. Brain Stimulation, 15(2), 391–402. 10.1016/j.brs.2022.02.006

46. Cavanagh, J. F., & Frank, M. J. (2014). Frontal theta as a mechanism for cognitive control. Trends in Cognitive Sciences, 18(8), 414–421. 10.1016/j.tics.2014.04.012

47. Ukharova, E., O’Meeghan, I., Granö, I., Sathyan, S., Iizuka, E., Tillander, C., Kainulainen, N., Alves, M. M., Engelhardt, M., Partanen, P., Vesterinen, M., Palva, S., Palva, J. M., Roine, T., Gogulski, J., Daskalakis, Z. J., Blumberger, D. M., Ilmoniemi, R. J., & Lioumis, P. (2026). Individualized TMS–EEG mapping reveals clinically relevant early prefrontal cortical activity in major depressive disorder. Clinical Neurophysiology, 2112427. 10.1016/j.clinph.2026.2112427

48. Biabani, M., Fornito, A., Goldsworthy, M., Thompson, S., Graetz, L., Semmler, J. G., Opie, G. M., Bellgrove, M. A., & Rogasch, N. C. (2024). Characterising the contribution of auditory and somatosensory inputs to TMS-evoked potentials following stimulation of prefrontal, premotor, and parietal cortex. Imaging Neuroscience, 2, imag–2–00349. 10.1162/imag_a_00349

49. Gordon, P. C., Desideri, D., Belardinelli, P., Zrenner, C., & Ziemann, U. (2018). Comparison of cortical EEG responses to realistic sham versus real TMS of human motor cortex. Brain Stimulation, 11(6), 1322–1330. 10.1016/j.brs.2018.08.003

50. Gordon, P. C., Jovellar, D. B., Song, Y., Zrenner, C., Belardinelli, P., Siebner, H. R., & Ziemann, U. (2021). Recording brain responses to TMS of primary motor cortex by EEG – utility of an optimized sham procedure. NeuroImage, 245, 118708. 10.1016/j.neuroimage.2021.118708

51. Fecchio, M., Pigorini, A., Comanducci, A., Sarasso, S., Casarotto, S., Premoli, I., Derchi, C.-C., Mazza, A., Russo, S., Resta, F., Ferrarelli, F., Mariotti, M., Ziemann, U., Massimini, M., & Rosanova, M. (2017). The spectral features of EEG responses to transcranial magnetic stimulation of the primary motor cortex depend on the amplitude of the motor evoked potentials. PLOS ONE, 12(9), e0184910. 10.1371/journal.pone.0184910

52. Petrichella, S., Johnson, N., & He, B. (2017). The influence of corticospinal activity on TMS-evoked activity and connectivity in healthy subjects: A TMS-EEG study. PLOS ONE, 12(4), e0174879. 10.1371/journal.pone.0174879

53. Cohen, P. (1988). Are statistics necessary? Biological Psychiatry, 23(1), 1–2. 10.1016/0006-3223(88)90100-X

54. Kerwin, L. J., Keller, C. J., Wu, W., Narayan, M., & Etkin, A. (2018). Test-retest reliability of transcranial magnetic stimulation EEG evoked potentials. Brain Stimulation, 11(3), 536–544. 10.1016/j.brs.2017.12.010

55. Benjamini, Y., & Hochberg, Y. (1995). Controlling the False Discovery Rate: A Practical and Powerful Approach to Multiple Testing. Journal of the Royal Statistical Society Series B: Statistical Methodology, 57(1), 289–300. 10.1111/j.2517-6161.1995.tb02031.x

56. Anderson, M. J., & Robinson, J. (2001). Permutation Tests for Linear Models. Australian & New Zealand Journal of Statistics, 43(1), 75–88. 10.1111/1467-842X.00156

57. Massimini, M., Peterchev, A. V., Vlachos, A., & Ziemann, U. (2026). Mechanisms of transcranial magnetic brain stimulation. Physiological Reviews, 106(4), 2267–2316. 10.1152/physrev.00026.2025

58. Salvatore, S. V., Lambert, P. M., Benz, A., Rensing, N. R., Wong, M., Zorumski, C. F., & Mennerick, S. (2024). Periodic and aperiodic changes to cortical EEG in response to pharmacological manipulation. Journal of Neurophysiology, 131(3), 529–540. 10.1152/jn.00445.2023

59. Hutt, A., McCulloch, M., Hudetz, A. G., Pariz, A., & Lefebvre, J. (2026). Noise-like fluctuations drive shifts in neuronal timescales and 1/f power spectra across brain states. PLOS Complex Systems, 3(9), e0000133. 10.1371/journal.pcsy.0000133

60. Belardinelli, P., König, F., Liang, C., Premoli, I., Desideri, D., Müller-Dahlhaus, F., Gordon, P. C., Zipser, C., Zrenner, C., & Ziemann, U. (2021). TMS-EEG signatures of glutamatergic neurotransmission in human cortex. Scientific Reports, 11(1), 8159. 10.1038/s41598-021-87533-z

61. Premoli, I., Castellanos, N., Rivolta, D., Belardinelli, P., Bajo, R., Zipser, C., Espenhahn, S., Heidegger, T., Müller-Dahlhaus, F., & Ziemann, U. (2014). TMS-EEG Signatures of GABAergic Neurotransmission in the Human Cortex. The Journal of Neuroscience, 34(16), 5603–5612. 10.1523/JNEUROSCI.5089-13.2014

62. Darmani, G., Zipser, C. M., Böhmer, G. M., Deschet, K., Müller-Dahlhaus, F., Belardinelli, P., Schwab, M., & Ziemann, U. (2016). Effects of the Selective α5-GABAAR Antagonist S44819 on Excitability in the Human Brain: A TMS–EMG and TMS–EEG Phase I Study. The Journal of Neuroscience, 36(49), 12312–12320. 10.1523/JNEUROSCI.1689-16.2016

63. Klimesch, W., Sauseng, P., & Hanslmayr, S. (2007). EEG alpha oscillations: The inhibition–timing hypothesis. Brain Research Reviews, 53(1), 63–88. 10.1016/j.brainresrev.2006.06.003

64. Peters, J. C., Reithler, J., Graaf, T. A. D., Schuhmann, T., Goebel, R., & Sack, A. T. (2020). Concurrent human TMS-EEG-fMRI enables monitoring of oscillatory brain state-dependent gating of cortico-subcortical network activity. Communications Biology, 3(1), 40. 10.1038/s42003-020-0764-0

65. Premoli, I., Bergmann, T. O., Fecchio, M., Rosanova, M., Biondi, A., Belardinelli, P., & Ziemann, U. (2017). The impact of GABAergic drugs on TMS-induced brain oscillations in human motor cortex. NeuroImage, 163, 1–12. 10.1016/j.neuroimage.2017.09.023

66. Rossiter, H. E., Davis, E. M., Clark, E. V., Boudrias, M.-H., & Ward, N. S. (2014). Beta oscillations reflect changes in motor cortex inhibition in healthy ageing. NeuroImage, 91, 360–365. 10.1016/j.neuroimage.2014.01.012

67. Poorganji, M., Zomorrodi, R., Zrenner, C., Bansal, A., Hawco, C., Hill, A. T., Hadas, I., Rajji, T. K., Chen, R., Zrenner, B., Voineskos, D., Blumberger, D. M., & Daskalakis, Z. J. (2023). Pre-Stimulus Power but Not Phase Predicts Prefrontal Cortical Excitability in TMS-EEG. Biosensors, 13(2), 220. 10.3390/bios13020220

68. Bergmann, T. O. (2018). Brain State-Dependent Brain Stimulation. Frontiers in Psychology, 9, 2108. 10.3389/fpsyg.2018.02108

69. Zrenner, C., Belardinelli, P., & Ziemann, U. (2026). Oscillatory brain state-dependent stimulation with transcranial magnetic stimulation combined with electroencephalography. Nature Protocols, 21(8), 3453–3490. 10.1038/s41596-025-01309-7

70. Van Hattem, T., Hougland, J. R., Ahola, O., Goetz, S. M., Humaidan, D., Jooß, A., & Ziemann, U. (2026). Brain-State-Resolved Consistency of Corticospinal Responses with EEG–TMS. bioRxiv. 10.64898/2026.05.25.727597

71. Baur, D., Galevska, D., Hussain, S., Cohen, L. G., Ziemann, U., & Zrenner, C. (2020). Induction of LTD-like corticospinal plasticity by low-frequency rTMS depends on pre-stimulus phase of sensorimotor μ-rhythm. Brain Stimulation, 13(6), 1580– 1587. 10.1016/j.brs.2020.09.005

72. Baur, D., Ermolova, M., Souza, V. H., Zrenner, C., & Ziemann, U. (2022). Phase-amplitude coupling in high-gamma frequency range induces LTP-like plasticity in human motor cortex: EEG-TMS evidence. Brain Stimulation, 15(6), 1508–1510. 10.1016/j.brs.2022.11.003

73. Kirchhoff, M., Humaidan, D., & Ziemann, U. (2024). *Closed-Loop phase selection in EEG-TMS using Bayesian Optimization* (arXiv:2410.05747). arXiv. 10.48550/arXiv.2410.05747

