## Supplementary Materials for "Aperiodic neural activity predicts cortical excitability in TMS-EEG"

**S.1 EEG preprocessing**

The TMS pulse artifact was removed from the post-stimulus window from –5 to 8 ms around the TMS pulse, and missing data was cubically interpolated. Noisy channels in the pre-stimulus or post-stimulus window were rejected from both windows using an automated algorithm based on global noise levels (standard deviation z-scores exceeding ± 2.5 for pre-stimulus EEG and ± 3 for post-stimulus EEG) and local peaks (mean ± SD rejected channels: M1 = 8.35% ± 5.31%; DLPFC = 10.65% ± 3.18%) [1]. Adjusted thresholds during channel rejection were applied for frontal channels to account for larger ocular artifacts (± 0.75). Independent component analysis (ICA) was then used to suppress ocular artifacts [2,3]. ICA filters fitted on the pre-stimulus window were removed from both the pre-stimulus and post-stimulus window. Pre-stimulus data were bandpass (2–80 Hz) and bandstop (48–52 Hz) filtered using a 4th-order zero-phase Butterworth filter. The pre-stimulus EEG was then re-referenced to the average reference and down-sampled to 1 kHz. Bad channels were reconstructed using spherical interpolation. Pre-stimulus EEG epochs were trimmed to –1.045 to –0.040 s to minimize edge artifacts and temporal smearing introduced by filtering. Bad trials were rejected using an automated rejection algorithm based on standard deviation z-scores within the pre-stimulus period exceeding ± 1.5 (mean ± SD rejected trials: M1 = 7.93% ± 4.85%; DLPFC = 3.62% ± 3.66%) [1]. The source-estimate-utilizing noise-discarding (SOUND) algorithm was used on the post-stimulus window to suppress external noise (e.g., electrode noise) [4]. TMS-evoked muscle artifacts were suppressed using the signal-space-projection-source-informed reconstruction (SSP–SIR) algorithm [5]. The post-stimulus window was baseline corrected using a window from –20 to –10 ms. Bad trials from the post-stimulus window were then automatically rejected using a standard deviation z-score threshold of ± 2.5 (mean ± SD rejected trials: M1 = 2.07% ± 0.50%; DLPFC = 2.31% ± 0.33%). The post-stimulus window was finally cropped to –0.020 to 0.5 s around the TMS pulse. The post-stimulus window was re-referenced to the average reference. For further details on the preprocessing pipeline, please refer to [1].

**S.2 Effect of periodic theta power on EL-TEP amplitude in DLPFC**

**S.2.1 Rationale & Methods**

Prefrontal theta oscillations are involved in cognitive processing and information transfer, as well as modulation of prefrontal cortical excitability [6]. Theta is sometimes used as a target EEG feature in brain-state-dependent TMS of the prefrontal cortex [7,8]. We evaluated the association between periodic theta power (mean spectral power between 4-8 Hz from the periodic component of the pre-stimulus EEG PSD) and EL-TEP amplitude in the DLPFC using the same methods as described in the main analysis.

**S.2.2 Results**

No significant differences were found in pre-stimulus theta power between the high- and low-response groups (High-response = –0.01 ± 0.03; Low-response = 0.003 ± 0.02; t(10) = 0.232, p = 0.821; Supplementary Figure 4A). At the single-trial level, there was a significant negative effect of theta on EL-TEP peak-to-peak amplitude (β = –0.020, SE = 0.004, t(9865) = –4.68, p < 0.001; Supplementary Figure 4B).

**S.3 Effects of aperiodic and periodic neural activity on reafferent somatosensory activity (N100-P200)**

**S.3.1 Rationale & Methods**

The EL-TEP comprises the early post-stimulus time window ~15–65 ms after TMS, which is thought to primarily reflect local cortical excitability, whereas later TEP components (> ~100 ms) are increasingly influenced by somatosensory-evoked activity [9–11]. As a control analysis, we quantified the effect of pre-stimulus aperiodic and periodic features on the N100-P200 peak-to-peak amplitude. The N100-P200 peak-to-peak amplitudes were extracted using the same procedures as for the EL-TEP time window. The final N100-P200 time window was 93–212 ms for M1 and 105–190 ms for DLPFC.

**S.3.2 Results**

For M1, no significant differences were found in pre-stimulus aperiodic exponent (High-response = 0.01 ± 0.02; Low-response = 0.008 ± 0.02; t(28) = 0.07, p = 0.943; Supplementary Figure 8C), alpha power (High-response = 0.02 ± 0.02; Low-response = –0.004 ± 0.01; t(28) = 0.92, p = 0.551; Supplementary Figure 8D), and beta power (High-response = 0.04 ± 0.02; Low-response = –0.02 ± 0.01; t(28) = 2.28, p = 0.092; Supplementary Figure 8E) between the high and low N100-P200 response groups. At the single-trial level, there was also no significant effect of aperiodic exponent (β = –0.0008, SE = 0.006, t(31280) = –0.125, p = 0.900; Supplementary Figure 8F), alpha power (β = –0.003, SE = 0.006, t(31280) = 0.415, p = 0.678; Supplementary Figure 8G), and beta power (β = 0.011, SE = 0.006, t(31280) = 1.80, p = 0.072; Supplementary Figure 8H) on N100-P200 peak-to-peak amplitude.

For DLPFC, no significant differences were found in pre-stimulus aperiodic exponent (High-response = –0.05 ± 0.03; Low-response = 0.03 ± 0.03; t(10) = –1.70, p = 0.179; Supplementary Figure 9C), alpha power (High-response = 0.01 ± 0.04; Low-response = –0.008 ± 0.03; t(10) = 0.42, p = 0.685; Supplementary Figure 9D) and beta power (High-response = –0.03 ± 0.03; Low-response = 0.03 ± 0.02; t(10) = –2.29, p = 0.135; Supplementary Figure 9E) between the high and low N100-P200 response groups. At the single-trial level, there was also no significant effect of aperiodic exponent (β = –0.016, SE = 0.011, t(9866) = –1.52, p = 0.129; Supplementary Figure 9F), alpha power (β = –0.003, SE = 0.011, t(9866) = –0.24, p = 0.811; Supplementary Figure 9G), and beta power (β = 0.004, SE = 0.011, t(9866) = 0.36, p = 0.719; Supplementary Figure 9H) on N100-P200 peak-to-peak amplitude.

**S.4 Earlier EL-TEP time window for M1**

**S.4.1 Rationale & Methods**

The M1 P60 component may be partly influenced by reafferent somatosensory feedback resulting from corticospinal activation with suprathreshold stimulation [12,13]. M1 was stimulated at 110% of resting motor threshold in this dataset. We therefore performed a control analysis at sensor level using a restricted EL-TEP window encompassing only approximately the P30–N45 complex (EL-TEP time window = 31–48 ms).

**S.4.2 Results**

Using an earlier EL-TEP time window, larger EL-TEP amplitude was associated with smaller aperiodic exponent (High-response = –0.15 ± 0.04; Low-response = 0.17 ± 0.02; t(28) = –5.20, p < 0.001; Supplementary Figure 3C) and lower beta power (High-response = –0.04 ± 0.03; Low-response = 0.07 ± 0.02; t(28) = –2.48, p = 0.029; Supplementary Figure 3E). No difference was found in pre-stimulus alpha power between response groups (High-response = –0.02 ± 0.03; Low-response = 0.009 ± 0.02; t(28) = –0.53, p = 0.600; Supplementary Figure 3D).

At the single-trial, aperiodic exponent (β = –0.051, SE = 0.003, t(31280) = –19.93, p < 0.001; Supplementary Figure 3F), alpha power (β = –0.015, SE = 0.003, t(31280) = –5.51, p < 0.001; Supplementary Figure 3G), and beta power (β = –0.008, SE = 0.003, t(31280) = –3.16, p < 0.001; Supplementary Figure 3H) were all negatively associated with EL-TEP amplitude.

**S.5 EEG source reconstruction**

**S.5.1 Rationale & Methods**

In addition to the primary sensor-space analyses, source-reconstructed activity was analyzed to help mitigate the influence of sensor-level non-neural noise sources and assess the cortical origin of the observed sensor effects.

T1-weighted MRIs were obtained with a 3T Siemens Prisma scanner and a 64-channel head coil (repetition time (TR) = 2.4 ms, echo time (TE) = 2.22 ms, slice thickness = 0.80 mm, and field of view (FoV) = 256 mm). Individual headmodels were created from the anatomical T1-weighted images using a three-shell Boundary Element Model (BEM) with customized scripts involving the use of Fieldtrip [14], Freesurfer [15], and Human Connectome Workbench packages. The source space was created for each participant with a cortical sheet model constructed based on the geometrical average of the boundaries between white and grey matter. A mesh of 15,684 vertices for each participant was obtained by adjusting the mesh points to a common spherical template for vertex-wise brain activation comparison across participants and downsampled. EEG channel locations from the experimental session were manually aligned to the scalp mesh using anatomical fiducials with the MATLAB Fieldtrip toolbox [14]. Leadfields were calculated for each participant and re-referenced to the average reference. Source activity was reconstructed using a linearly constrained minimum variance (LCMV) beamformer, and the covariance matrix to create the filter used Tikhonov regularization with a parameter set to 10 times the maximum sensor power [16,17].

For M1, we defined a custom hand-knob ROI based on the intersection of two macro-parcels [18]: the union of three sensorimotor Glasser parcels (BA4 (M1), S1, and Area 1) [19] and the union of six Brodmann areas for the left hand (BA Hand 1-6). For DLPFC, five ROIs were selected based on the Glasser parcellation atlas (8Ad, 8Av, 8C, 46, and p9-46v) [19]. Source signals were averaged across the ROIs for both pre- and post-window EEG. Pre-stimulus feature extraction and EL-TEP amplitude calculation from the source signals were done the same as for the sensor-based analysis (see Section *2.5.3 Post-stimulus window: peak-to-peak EL-TEP amplitudes*.) The final EL-TEP time window in source space was 30–62 ms for M1 and 20–64 ms for DLPFC.

**S.5.2 Results**

In concordance with the sensor-level outcomes, larger source EL-TEPs in M1 were associated with smaller pre-stimulus aperiodic exponent (High-response = –0.17 ± 0.04; Low-response = 0.18 ± 0.02; t(28) = –5.82, p < 0.001; Supplementary Figure 5C) and lower beta power (High-response = –0.08 ± 0.02; Low-response = 0.08 ± 0.02; t(28) = –4.03, p < 0.001; Supplementary Figure 5E), but there was no difference in pre-stimulus alpha power between response groups (High-response = –0.01 ± 0.02; Low-response = –0.02 ± 0.02; t(28) = 0.12, p = 0.902; Supplementary Figure 5D).

Similar to the sensor-level findings, larger source EL-TEPs in DLPFC were associated with smaller pre-stimulus aperiodic exponent (High-response = –0.23 ± 0.08; Low-response = 0.22 ± 0.05; t(10) = –4.72, p = 0.002; Supplementary Figure 6C). No differences between response groups were found in alpha power (High-response = 0.09 ± 0.05; Low-response = –0.10 ± 0.04; t(10) = 2.20, p = 0.078; Supplementary Figure 6D) or beta power (High-response = 0.01 ± 0.04; Low-response = –0.05 ± 0.03; t(10) = 0.89, p = 0.392; Supplementary Figure 6E).

**S.6 Dummy triggers in resting-state EEG**

**S.6.1 Rationale & Methods**

To assess whether the observed associations between pre-stimulus aperiodic and periodic features and EL-TEP amplitude could be explained by ongoing EEG dynamics rather than TMS-related responses, we performed a control analysis at sensor level in which artificial trigger markers were inserted at random intervals of 2–2.5 s in 5-minute resting-state EEG. The resting-state EEG data were then analyzed using the same preprocessing and feature-extraction procedures used for the TMS-EEG data. We subsequently examined the relationship between pre-stimulus aperiodic and periodic EEG features and the amplitude of the corresponding post-stimulus EEG signal.

Because resting-state EEG does not contain a TMS-evoked response, post-trigger EEG peak-to-peak amplitudes primarily reflect ongoing EEG fluctuations and noise. To avoid selecting or grouping trials based on this and potentially fitting noise, trials were split based on the pre-stimulus EEG feature of interest rather than post-stimulus EEG signal amplitude, as was done in the main analysis. Trials were divided into tertiles based on the pre-stimulus aperiodic exponent, with the upper and lower tertiles used to compare epochs characterized by relatively higher versus lower exponent values. A tertile split was used because of the limited number of available trials. For the M1 data, an average of 111.2 trials (SD = 8.3) per participant were available for this analysis.

**S.6.2 Results**

For M1, there were no significant differences in post-stimulus EEG peak-to-peak amplitudes between high and low aperiodic exponent (High mean ± SEM = 0.28 ± 0.06; Low mean ± SEM = 0.34 ± 0.08; t(28) = –0.93, p-uncorrected = 0.361; Supplementary Figure 10A), alpha power (High mean ± SEM = 0.26 ± 0.06; Low mean ± SEM = 0.32 ± 0.07; t(28) = –0.97, p-uncorrected = 0.342; Supplementary Figure 10B), or beta power (High mean ± SEM = 0.32 ± 0.09; Low mean ± SEM = 0.29 ± 0.08; t(28) = 0.39, p-uncorrected = 0.698; Supplementary Figure 10C).

For DLPFC, there were also no significant differences in post-stimulus EEG peak-to-peak amplitudes between high and low aperiodic exponent (High mean ± SEM = 0.55 ± 0.14; Low mean ± SEM = 0.38 ± 0.13; t(10) = 1.08, p-uncorrected = 0.307; Supplementary Figure 10D), alpha power (High mean ± SEM = 0.61 ± 0.09; Low mean ± SEM = 0.42 ± 0.12; t(10) = 1.58, p-uncorrected = 0.145; Supplementary Figure 10E), or beta power (High mean ± SEM = 0.64 ± 0.05; Low mean ± SEM = 0.45 ± 0.10; t(10) = 1.88, p-uncorrected = 0.090; Supplementary Figure 10F).

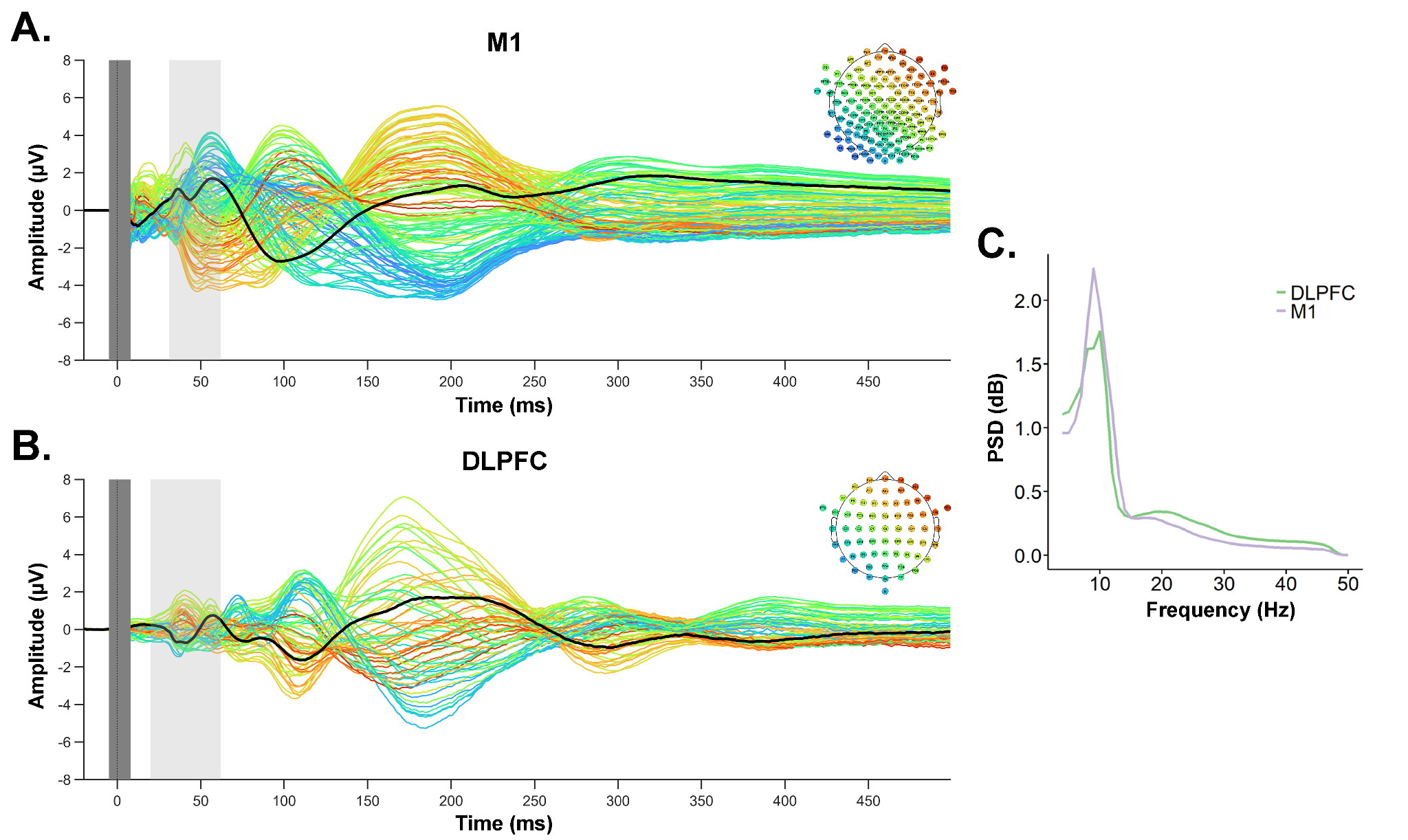

**Supplementary Figure 1. Sensor-level grand-average TMS-evoked EEG responses following M1 and DLPFC stimulation. (A)** Butterfly plots of TEPs following M1 stimulation recorded with a 126-channel EEG cap. **(B)** Butterfly plots of TEPs following DLPFC stimulation recorded with a 64-channel EEG cap. The vertical light gray window represents the EL-TEP time window (M1: 31-62 ms; DLPFC: 20-62 ms). The vertical dark gray window represents the data affected by the TMS pulse artefact that was removed and interpolated. The black line represents the ROI average waveform (M1 ROI: C3, C1, FC1, and FC3; DLPFC ROI: F3, F1, FC3, F5, FC1, and FC5). **(C)** Grand-average pre-stimulus PSD for M1 and DLPFC.

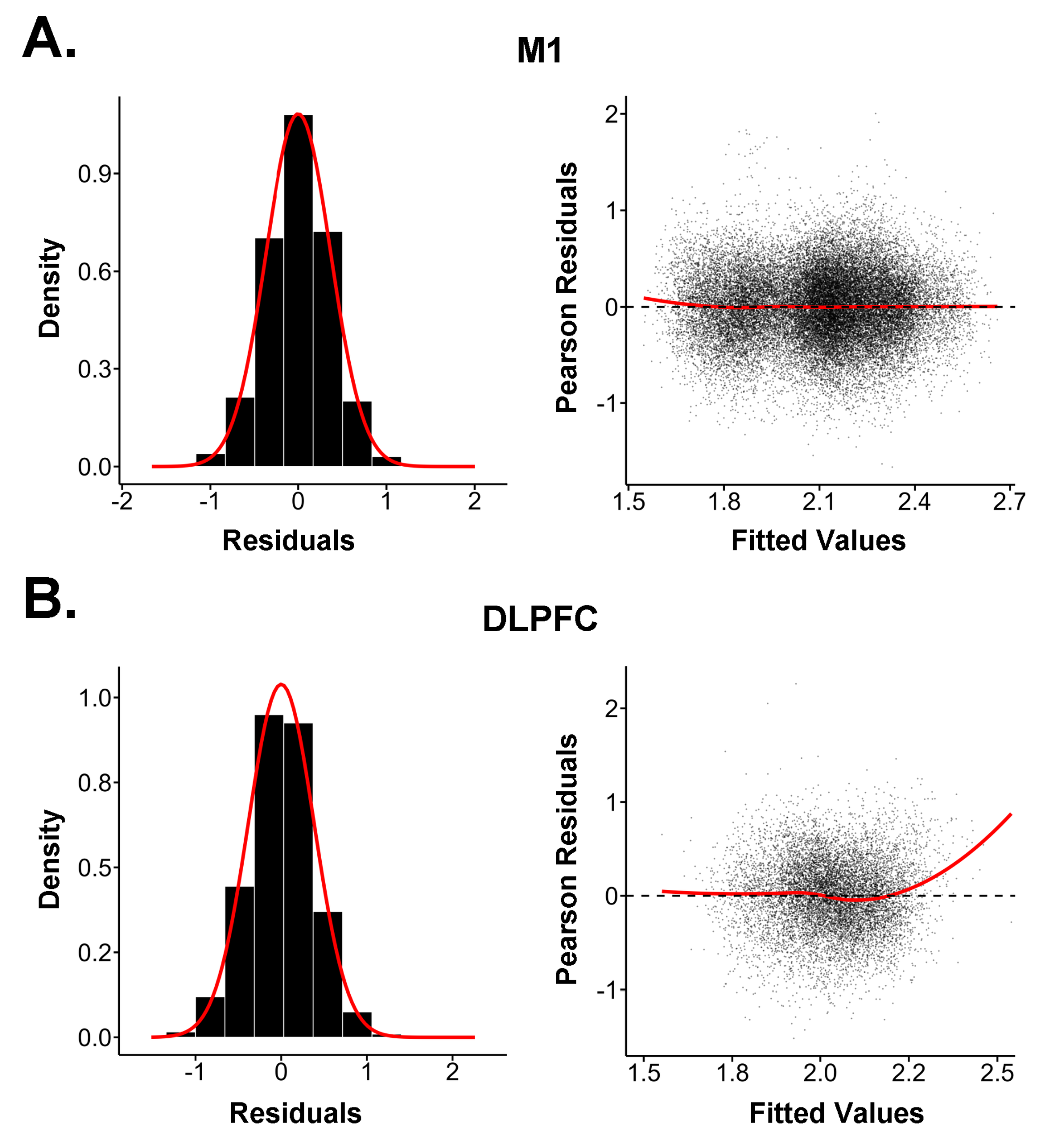

**Supplementary Figure 2. Model diagnostics for single-trial linear mixed-effects models**. Density distributions of the linear mixed-effects models residuals were approximately centered around zero which indicates that there was no systematic bias in the model predictions. Pearson residuals as a function of the fitted values show no major deviations from model assumptions in M1. The DLPFC model showed some deviations, indicating the model underpredicts higher fitted values. The red line reflects the loess regression fit.

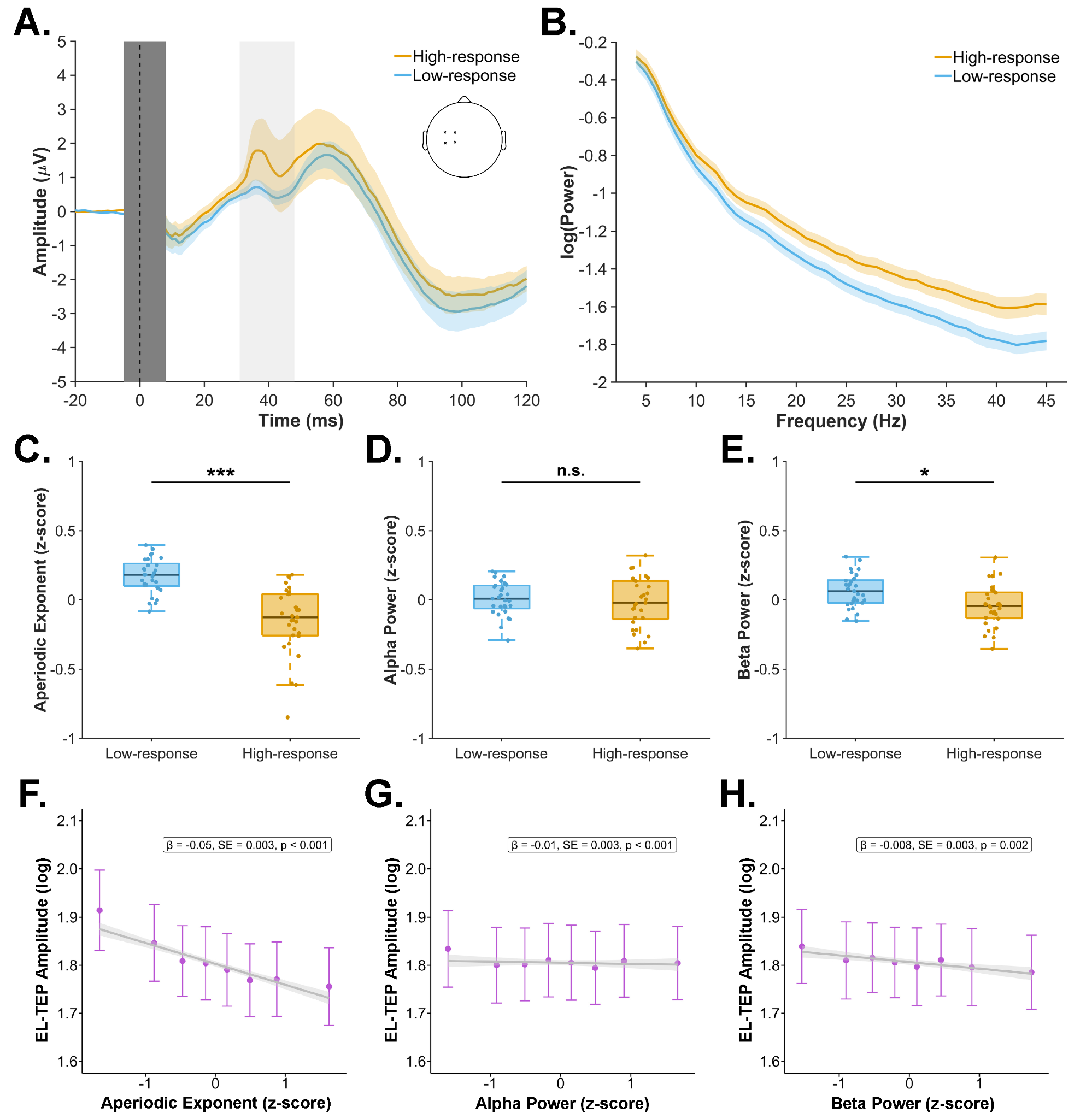

**Supplementary Figure 3. Effect of aperiodic and periodic EEG features on EL-TEP amplitude in M1 using an earlier EL-TEP time window (31–48 ms) to reduce the potential influence of reafferent feedback. (A)** Sensor-level grand-average waveforms across all participants across all participants (N = 29) for the high- and low-response groups, defined as the 150 highest and lowest EL-TEP trials, respectively. The ROI included electrodes C3, C1, FC1, and FC3, as indicated in the inset. The vertical light gray window represents the EL-TEP time window. The vertical dark gray window represents the data affected by the TMS pulse artifact that was removed and interpolated. The coloured shaded areas around the waveform show ± SEM. **(B)** Average aperiodic fits for the high- and low-response groups. The high-response group exhibited a flatter aperiodic slope (i.e., smaller aperiodic exponent) than the low-response group. The coloured shaded areas around the aperiodic fit show ± SEM. **(C–E)** Box plots show the median, interquartile range, and whiskers extending to 1.5 times the interquartile range of the pre-stimulus EEG feature for the high- and low-response groups. Y-axes show z-score normalized pre-stimulus EEG feature values. Dots represent data from individual participants. Compared with the low-response group, the high-response group showed a significantly smaller pre-stimulus aperiodic exponent and lower beta power. Pre-stimulus alpha power did not differ significantly between groups. * p < .05; *** p < .001. **(F–H)** Single-trial analyses revealed significant negative associations between EL-TEP amplitude and the aperiodic exponent, alpha power, and beta power. Grey lines represent the linear mixed-effects model-estimated relationship between each pre-stimulus EEG feature and EL-TEP amplitude. Grey shaded areas indicating 95% confidence intervals. Data points show binned mean values across octiles plotted as mean ± SEM. X-axes show z-score normalized pre-stimulus EEG feature values. Text boxes report the fixed-effect estimates (β), standard errors (SE), and p-values for each EEG feature.

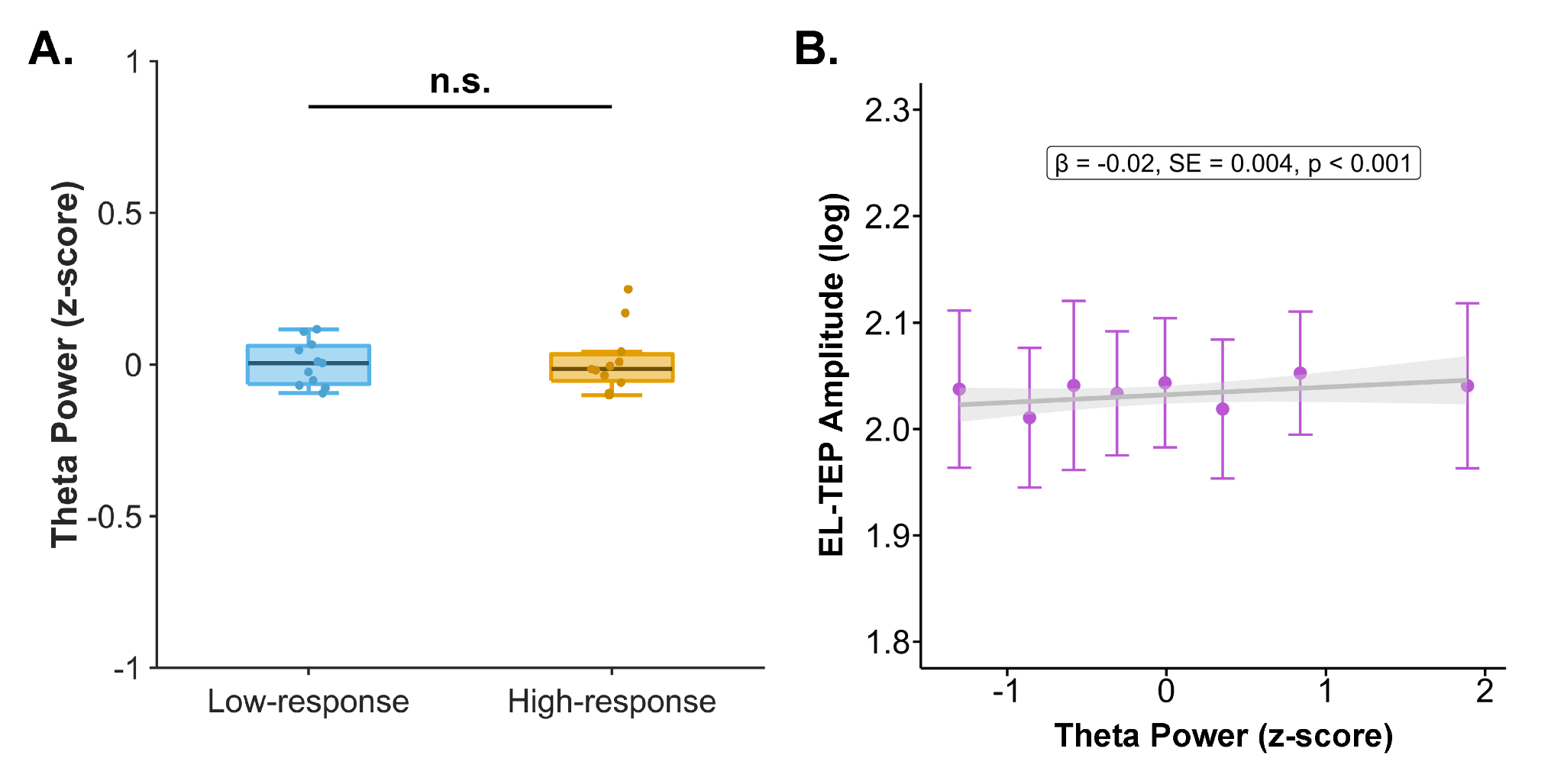

**Supplementary Figure 4. Effect of periodic theta power on EL-TEP amplitude in DLPFC. (A)** Box plots show the median, interquartile range, and whiskers extending to 1.5 times the interquartile range of the pre-stimulus theta power for the high- and low-response groups. Y-axes show z-score normalized pre-stimulus theta power values. Dots represent data from individual participants. There was no significant difference between the two groups. **(B)** Single-trial predictions of EL-TEP amplitude using pre-stimulus theta power in DLPFC. A linear mixed-effects model was fit to all trials to assess the effect of pre-stimulus theta power on the EL-TEP amplitude, with aperiodic exponent, theta power, alpha power, and beta power as fixed effects, and participant as a random intercept. Grey lines represent the linear mixed-effects model-estimated relationship between theta power and EL-TEP amplitude. Grey shaded areas indicating 95% confidence intervals. Data points show binned mean values across octiles plotted as mean ± SEM. X-axes show z-score normalized pre-stimulus EEG feature values. The text box reports the fixed-effect estimate (β), standard error (SE), and p-value.

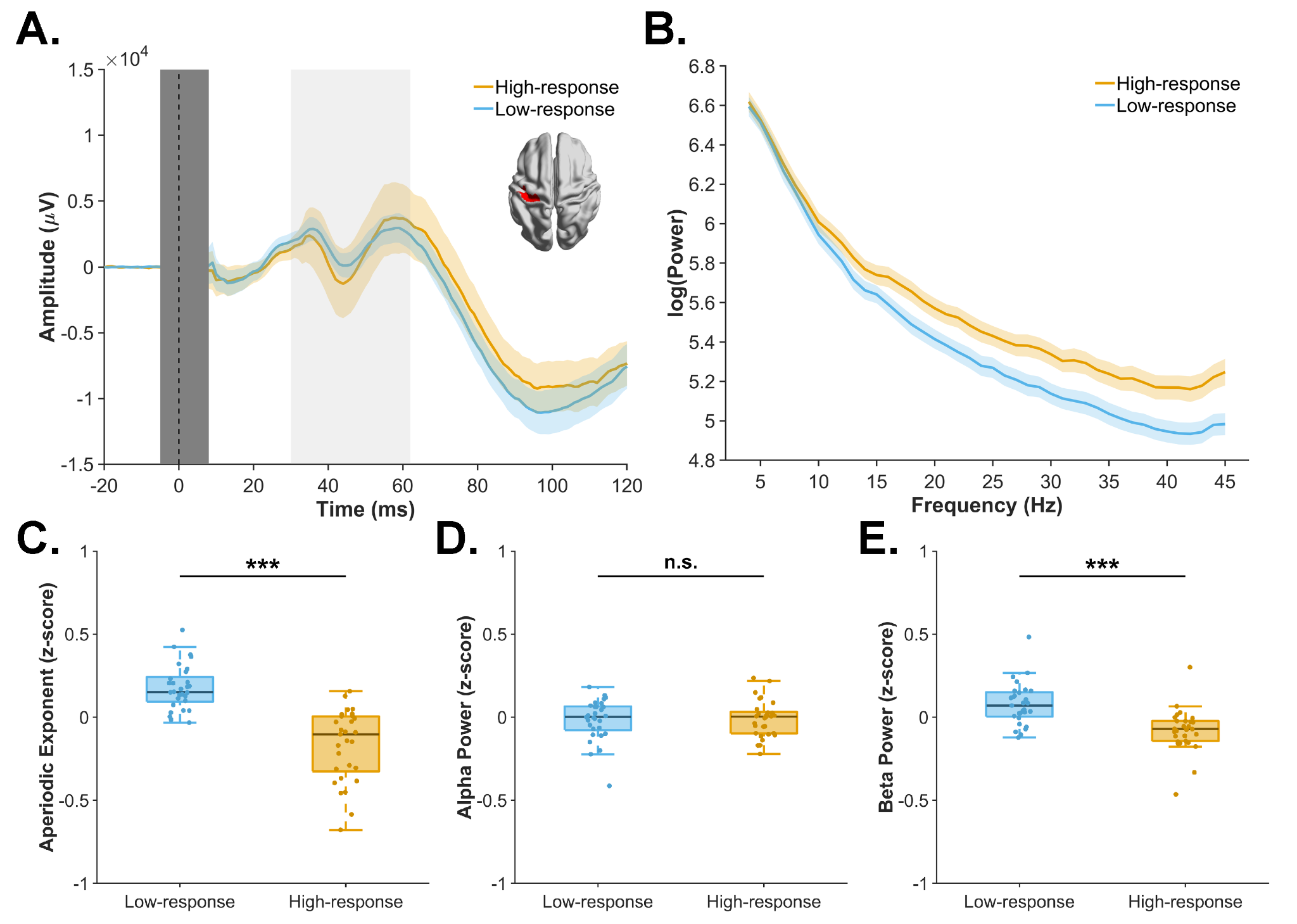

**Supplementary Figure 5. Effect of aperiodic and periodic EEG features on EL-TEP amplitude in M1 at the source level. (A)** Source-level grand-average waveforms across all participants (N = 29) for the high- and low-response groups, defined as the 150 highest and lowest EL-TEP trials, respectively. The ROI included a custom hand-knob ROI based on the intersection of two macro-parcels: the union of three sensorimotor Glasser parcels (BA4 (M1), S1, and Area 1) and the union of six Broadmann areas for the left hand (BA Hand 1-6), as indicated in the inset. The vertical light gray window represents the EL-TEP time window (31–62 ms). The vertical dark gray window represents the data affected by the TMS pulse artifact that was removed and interpolated. The coloured shaded areas around the waveform show ± SEM. **(B)** Average aperiodic fits for the high- and low-response groups. The high-response group exhibited a flatter aperiodic slope (i.e., smaller aperiodic exponent) than the low-response group. The coloured shaded areas around the aperiodic fit show ± SEM. **(C–E)** Box plots show the median, interquartile range, and whiskers extending to 1.5 times the interquartile range of the pre-stimulus EEG feature for the high- and low-response groups. Y-axes show z-score normalized pre-stimulus EEG feature values. Dots represent data from individual participants. Compared with the low-response group, the high-response group showed a significantly smaller pre-stimulus aperiodic exponent and lower beta power. Pre-stimulus alpha power was not significantly different between groups. *** p < .001.

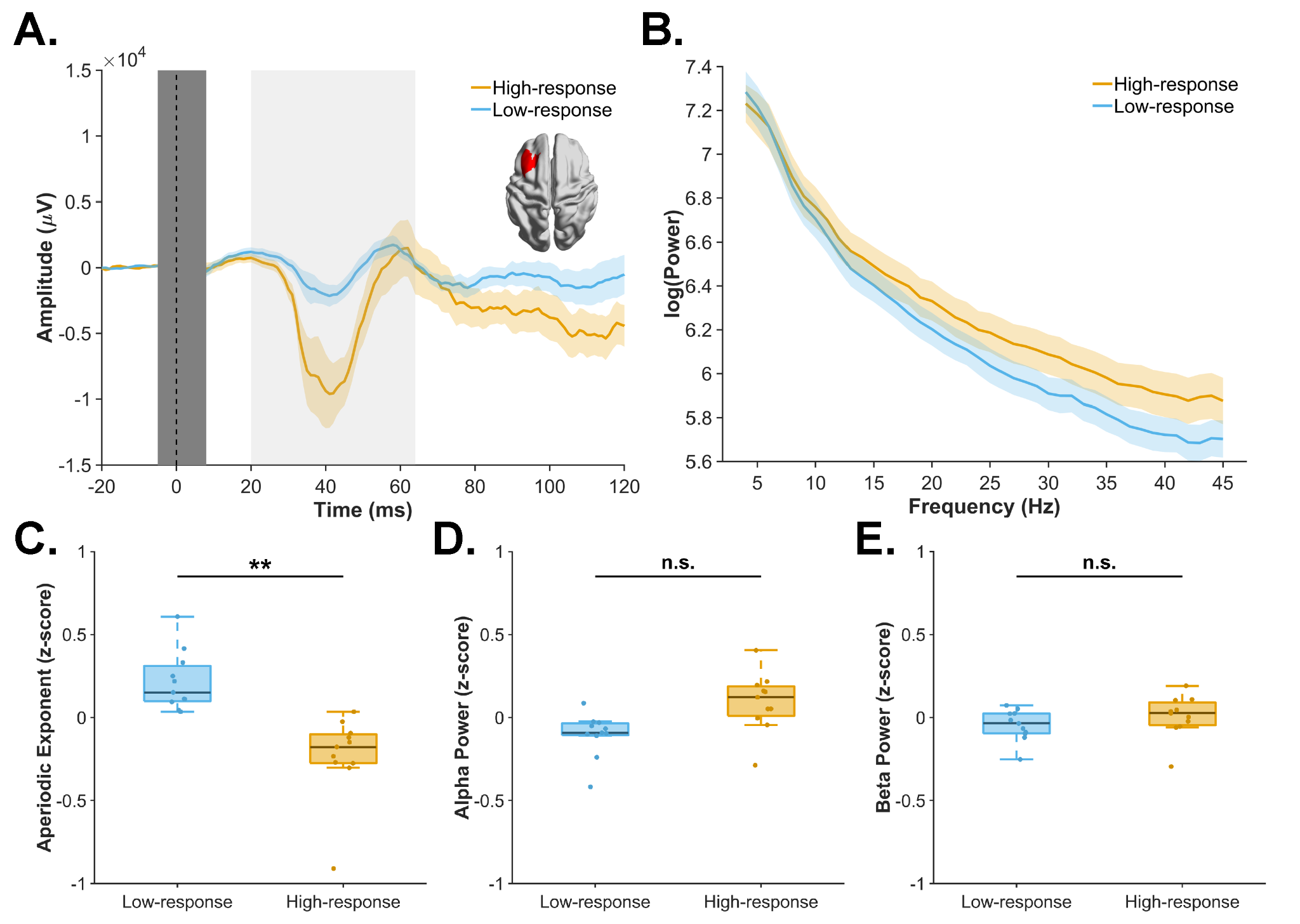

**Supplementary Figure 6. Effect of aperiodic and periodic EEG features on EL-TEP amplitude in DLPFC at the source level. (A)** Source-level grand-average waveforms across all participants (N = 11) for the high- and low-response groups, defined as the 150 highest and lowest EL-TEP trials, respectively. The ROI included Glasser atlas parcels 8Ad, 8Av, 8C, 46, and p9-46v, as indicated in the inset. The vertical light gray window represents the EL-TEP time window. The vertical dark gray window represents the data affected by the TMS pulse artifact that was removed and interpolated. The coloured shaded areas around the waveform show ± SEM. **(B)** Average aperiodic fits for the high- and low-response groups. The high-response group exhibited a flatter aperiodic slope (i.e., smaller aperiodic exponent) than the low-response group. The coloured shaded areas around the aperiodic fit show ± SEM. **(C–E)** Box plots show the median, interquartile range, and whiskers extending to 1.5 times the interquartile range of the pre-stimulus EEG feature for the high- and low-response groups. Y-axes show z-score normalized pre-stimulus EEG feature values. Dots represent data from individual participants. Compared with the low-response group, the high-response group showed a significantly smaller pre-stimulus aperiodic exponent. Pre-stimulus alpha and beta power did not differ significantly between groups. * p < .05.

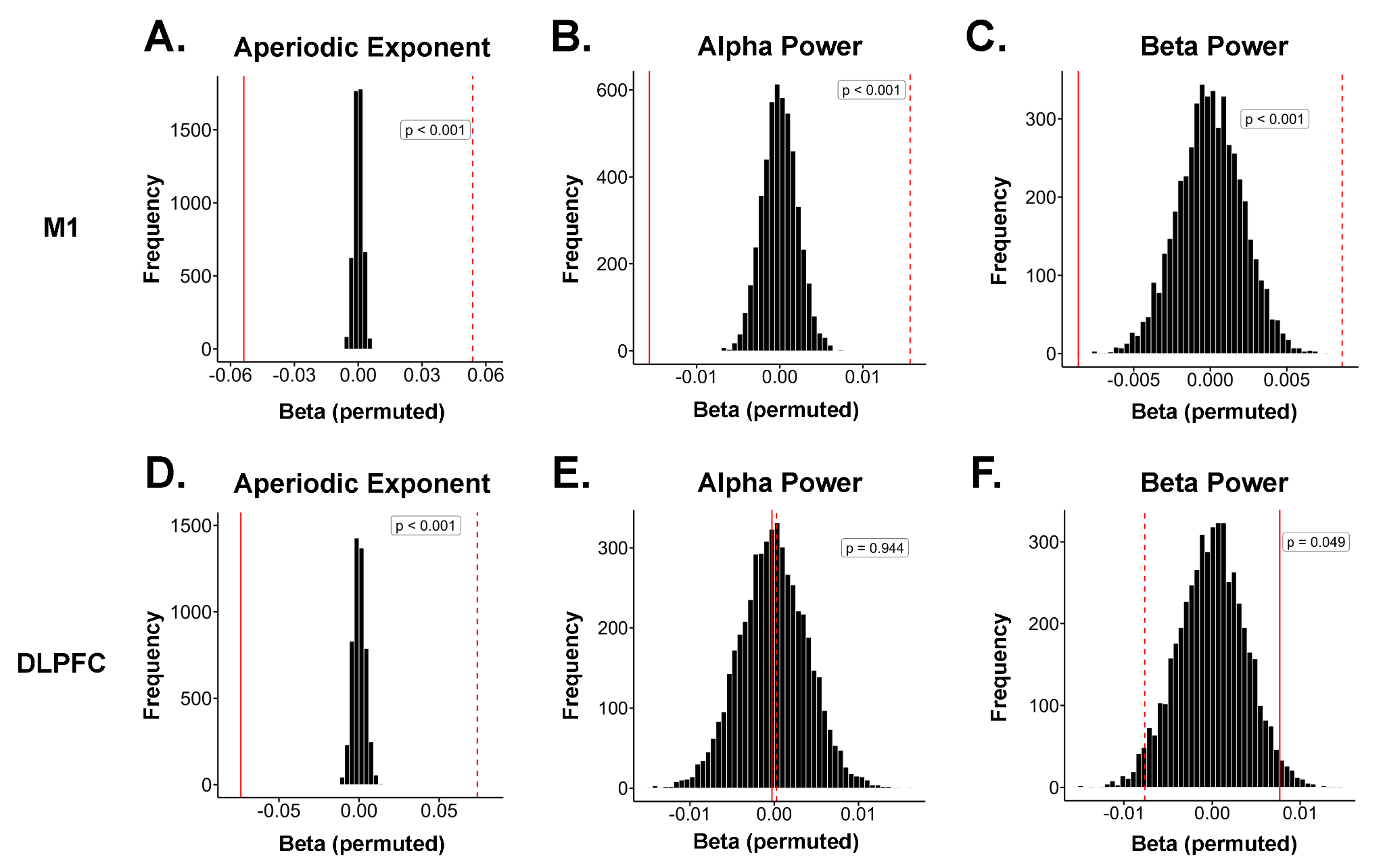

**Supplementary Figure 7. Permutation tests for effect of aperiodic and periodic EEG features on EL-TEP amplitude at sensor-level.** Null distributions of single-trial linear mixed-effects fixed-effect beta coefficients of the aperiodic exponent, alpha power, and beta power predictors were generated using 5000 permutations. The solid red line indicates the observed beta coefficient from the linear mixed-effects model. The dashed red line indicates the sign-reversed beta coefficient, reflecting the opposite tail of the two-tailed test. Permutation *p*-values reflect the proportion of permuted coefficients whose absolute value equaled or exceeded that of the original model. The results confirmed the statistical significance of all the original linear mixed-effects models, except for the association between beta power and EL-TEP amplitude in DLPFC, which narrowly exceeded chance-level expectations in the permutation test.

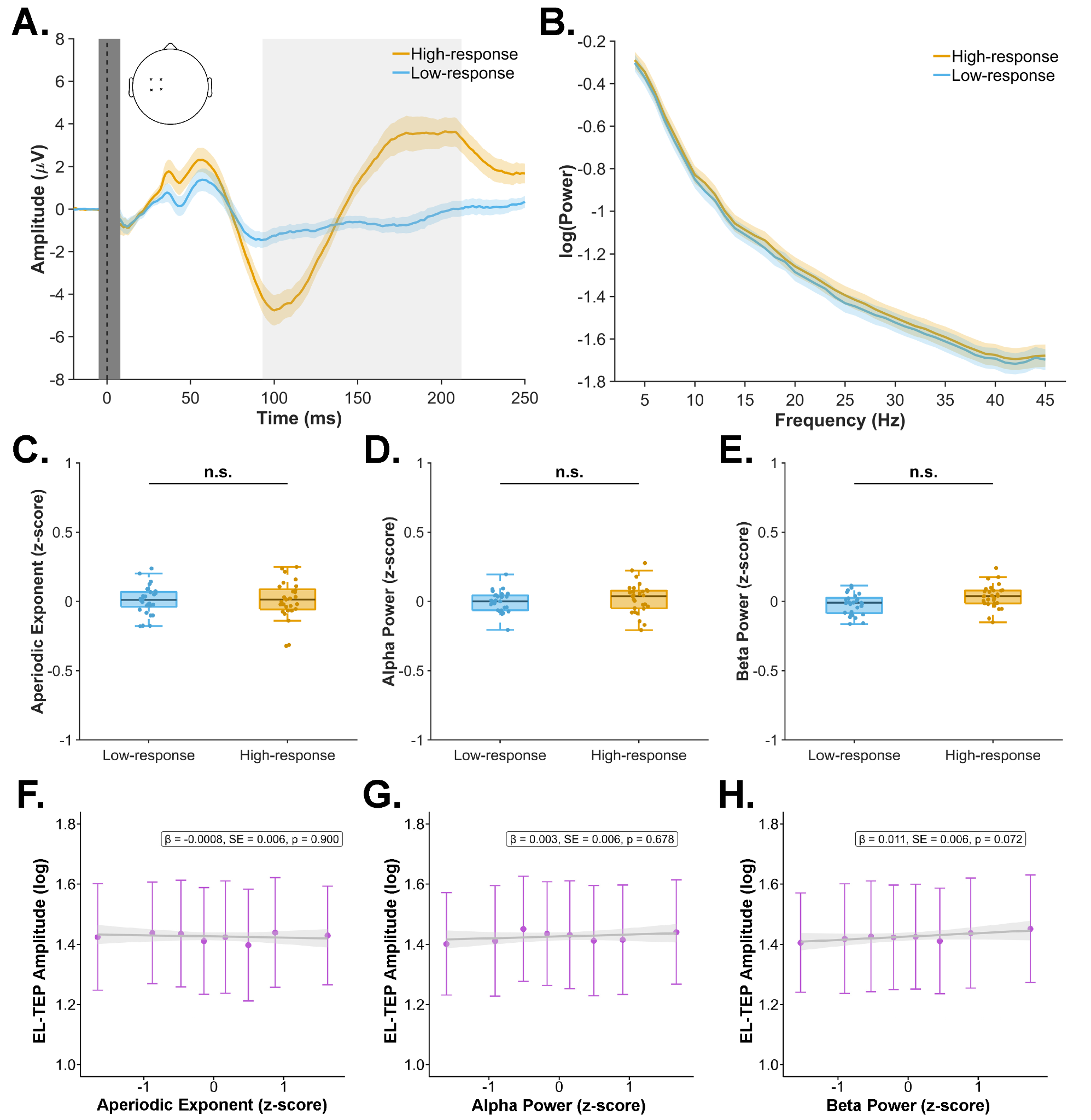

**Supplementary Figure 8. Effect of aperiodic and periodic EEG features on reafferent somatosensory activity (N100-P200) in M1. (A)** Sensor-level grand-average waveforms across all participants across all participants (N = 29) for the high- and low-response groups, defined as the 150 highest and lowest N100-P200 peak-to-peak amplitudes, respectively. The ROI included electrodes C3, C1, FC1, and FC3, as indicated in the inset. The vertical light gray window represents the N100-P200 time window. The vertical dark gray window represents the data affected by the TMS pulse artifact that was removed and interpolated. The coloured shaded areas around the waveform show ± SEM. **(B)** Average aperiodic fits for the high- and low-response groups. The high- and low-response groups exhibited similar aperiodic slopes (i.e., aperiodic exponent). The coloured shaded areas around the aperiodic fit show ± SEM. **(C–E)** Box plots show the median, interquartile range, and whiskers extending to 1.5 times the interquartile range of the pre-stimulus EEG feature for the high- and low-response groups. Y-axes show z-score normalized pre-stimulus EEG feature values. Dots represent data from individual participants. Pre-stimulus aperiodic exponent, alpha and beta power did not differ significantly between groups. **(F–H)** Single-trial analyses did not reveal significant associations between N100-P200 amplitude and the aperiodic exponent, alpha power, and beta power. Grey lines represent the linear mixed-effects model-estimated relationship between each pre-stimulus EEG feature and EL-TEP amplitude. Grey shaded areas indicating 95% confidence intervals. Data points show binned mean values across octiles plotted as mean ± SEM. X-axes show z-score normalized pre-stimulus EEG feature values. Text boxes report the fixed-effect estimates (β), standard errors (SE), and p-values for each EEG feature.

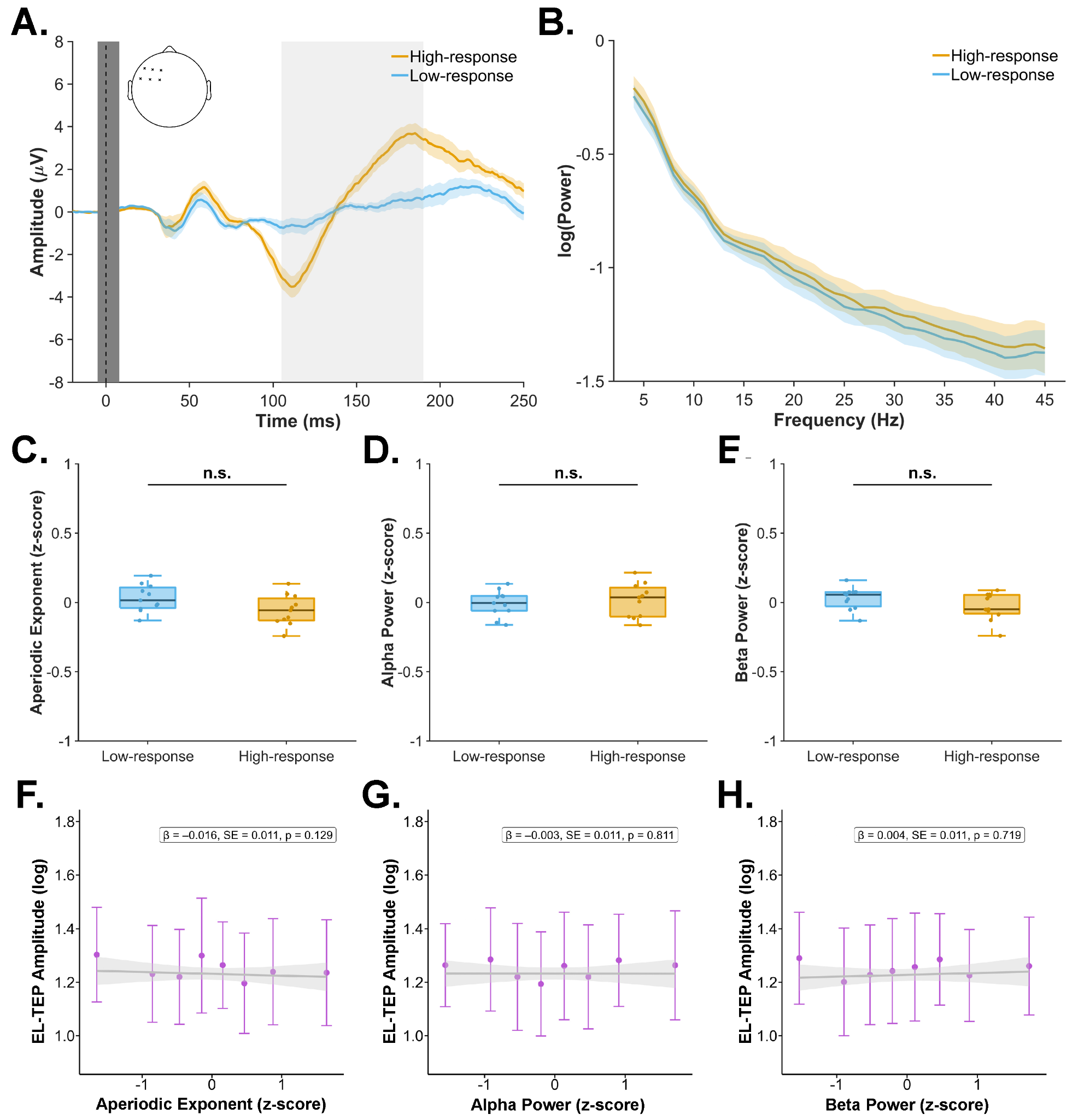

**Supplementary Figure 9. Effect of aperiodic and periodic EEG features on reafferent somatosensory activity (N100-P200) in DLPFC. (A)** Sensor-level grand-average waveforms across all participants across all participants (N = 11) for the high- and low-response groups, defined as the 150 highest and lowest N100-P200 peak-to-peak amplitudes, respectively. The ROI included electrodes F3, F1, FC3, F5, FC1, and FC5, as indicated in the inset. The vertical light gray window represents the N100-P200 time window. The vertical dark gray window represents the data affected by the TMS pulse artifact that was removed and interpolated. The coloured shaded areas around the waveform show ± SEM. **(B)** Average aperiodic fits for the high- and low-response groups. The high- and low-response groups exhibited similar aperiodic slopes (i.e., aperiodic exponent). The coloured shaded areas around the aperiodic fit show ± SEM. **(C–E)** Box plots show the median, interquartile range, and whiskers extending to 1.5 times the interquartile range of the pre-stimulus EEG feature for the high- and low-response groups. Y-axes show z-score normalized pre-stimulus EEG feature values. Dots represent data from individual participants. Pre-stimulus aperiodic exponent, alpha and beta power did not differ significantly between groups. **(F–H)** Single-trial analyses did not reveal significant associations between N100-P200 amplitude and the aperiodic exponent, alpha power, and beta power. Grey lines represent the linear mixed-effects model-estimated relationship between each pre-stimulus EEG feature and EL-TEP amplitude. Grey shaded areas indicating 95% confidence intervals. Data points show binned mean values across octiles plotted as mean ± SEM. X-axes show z-score normalized pre-stimulus EEG feature values. Text boxes report the fixed-effect estimates (β), standard errors (SE), and p-values for each EEG feature.

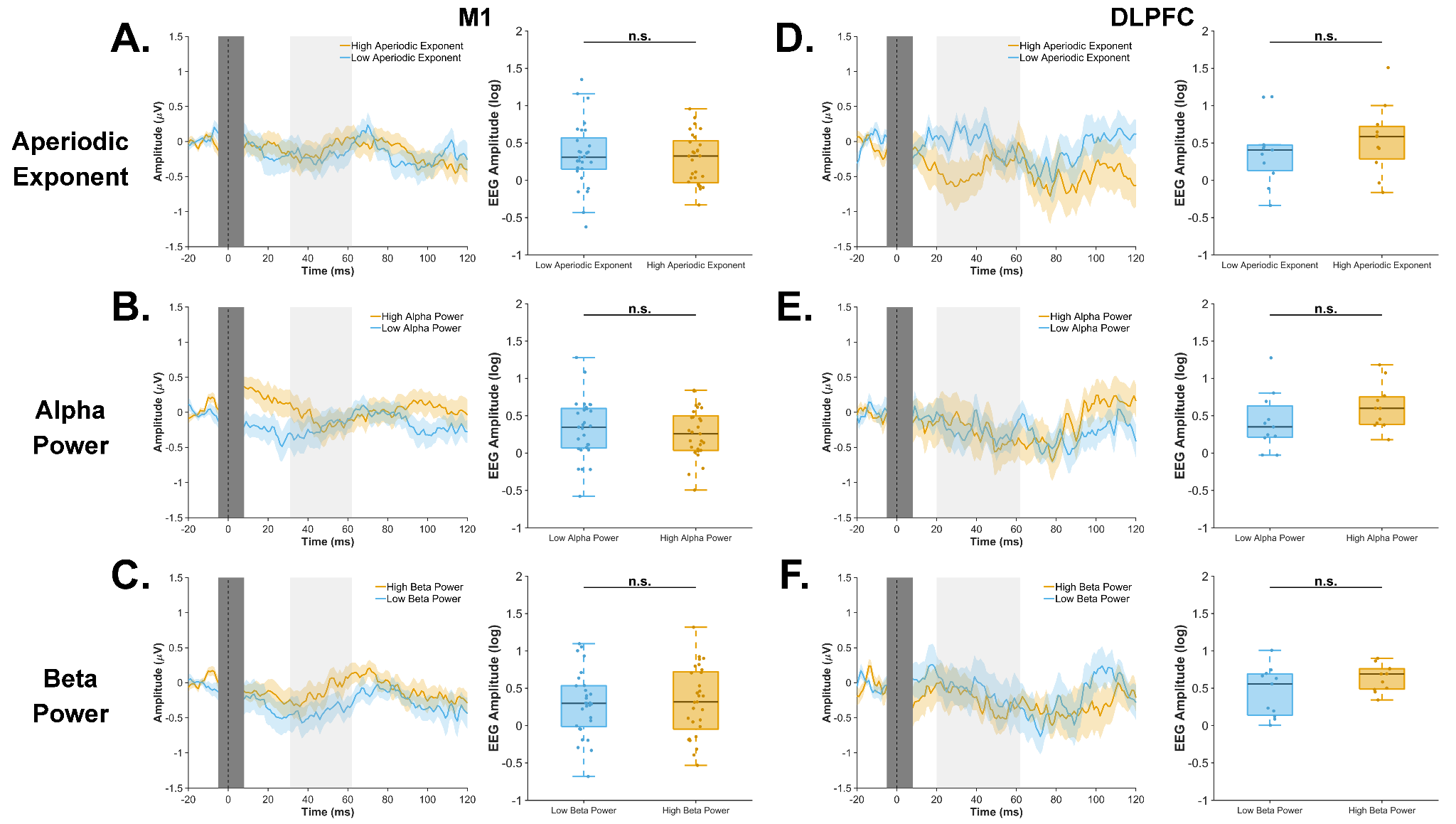

**Supplementary Figure 10. Dummy triggers in resting-state EEG. (A-C)** Tertile split based on pre-stimulus aperiodic exponent, alpha power, and beta power in M1. **(D-F)** Tertile split based on pre-stimulus aperiodic exponent, alpha power, and beta power in DLPFC. The left subplot shows sensor-level grand-average waveforms across all participants for the high- and low-feature groups, defined as the third and first tertiles, respectively. The vertical light gray window represents the EL-TEP time window. The vertical dark gray window represents the data affected by the TMS pulse artifact that was removed and interpolated. The coloured shaded areas around the waveform show ± SEM. The right subplot shows box plots with the median, interquartile range, and whiskers extending to 1.5 times the interquartile range of the EEG amplitude for the high- and low-feature groups. Y-axes show log-transformed peak-to-peak EEG amplitudes. Dots represent data from individual participants.
**Supplementary References**

1.	Ahola, O., Haxel, L., Ermolova, M., Humaidan, D., Mutanen, T. P., Laine, M., Makkonen, M., Ukharova, E., Roine, T., Lioumis, P., Guidotti, R., Llmoniemi, R. J., & Ziemann, U. (2025). Predictive modeling of TMS-evoked responses: Unraveling instantaneous excitability states. *NeuroImage*, *322*, 121553. https://doi.org/10.1016/j.neuroimage.2025.121553

2.	Hyvärinen, A., & Oja, E. (2000). Independent component analysis: Algorithms and applications. *Neural Networks*, *13*(4–5), 411–430. https://doi.org/10.1016/S0893-6080(00)00026-5

3.	Pion-Tonachini, L., Kreutz-Delgado, K., & Makeig, S. (2019). ICLabel: An automated electroencephalographic independent component classifier, dataset, and website. *NeuroImage*, *198*, 181–197. https://doi.org/10.1016/j.neuroimage.2019.05.026

4.	Mutanen, T. P., Metsomaa, J., Liljander, S., & Ilmoniemi, R. J. (2018). Automatic and robust noise suppression in EEG and MEG: The SOUND algorithm. *NeuroImage*, *166*, 135–151. https://doi.org/10.1016/j.neuroimage.2017.10.021

5.	Mutanen, T. P., Kukkonen, M., Nieminen, J. O., Stenroos, M., Sarvas, J., & Ilmoniemi, R. J. (2016). Recovering TMS-evoked EEG responses masked by muscle artifacts. *NeuroImage*, *139*, 157–166. https://doi.org/10.1016/j.neuroimage.2016.05.028

6.	Cavanagh, J. F., & Frank, M. J. (2014). Frontal theta as a mechanism for cognitive control. *Trends in Cognitive Sciences*, *18*(8), 414–421. https://doi.org/10.1016/j.tics.2014.04.012

7.	Gordon, P. C., Dörre, S., Belardinelli, P., Stenroos, M., Zrenner, B., Ziemann, U., & Zrenner, C. (2021). Prefrontal Theta-Phase Synchronized Brain Stimulation With Real-Time EEG-Triggered TMS. *Frontiers in Human Neuroscience*, *15*, 691821. https://doi.org/10.3389/fnhum.2021.691821

8.	Gordon, P. C., Belardinelli, P., Stenroos, M., Ziemann, U., & Zrenner, C. (2022). Prefrontal theta phase-dependent rTMS-induced plasticity of cortical and behavioral responses in human cortex. *Brain Stimulation*, *15*(2), 391–402. https://doi.org/10.1016/j.brs.2022.02.006

9.	Biabani, M., Fornito, A., Goldsworthy, M., Thompson, S., Graetz, L., Semmler, J. G., Opie, G. M., Bellgrove, M. A., & Rogasch, N. C. (2024). Characterising the contribution of auditory and somatosensory inputs to TMS-evoked potentials following stimulation of prefrontal, premotor, and parietal cortex. *Imaging Neuroscience*, *2*, imag–2–00349. https://doi.org/10.1162/imag_a_00349

10.	Gordon, P. C., Desideri, D., Belardinelli, P., Zrenner, C., & Ziemann, U. (2018). Comparison of cortical EEG responses to realistic sham versus real TMS of human motor cortex. *Brain Stimulation*, *11*(6), 1322–1330. https://doi.org/10.1016/j.brs.2018.08.003

11.	Gordon, P. C., Jovellar, D. B., Song, Y., Zrenner, C., Belardinelli, P., Siebner, H. R., & Ziemann, U. (2021). Recording brain responses to TMS of primary motor cortex by EEG – utility of an optimized sham procedure. *NeuroImage*, *245*, 118708. https://doi.org/10.1016/j.neuroimage.2021.118708

12.	Fecchio, M., Pigorini, A., Comanducci, A., Sarasso, S., Casarotto, S., Premoli, I., Derchi, C.-C., Mazza, A., Russo, S., Resta, F., Ferrarelli, F., Mariotti, M., Ziemann, U., Massimini, M., & Rosanova, M. (2017). The spectral features of EEG responses to transcranial magnetic stimulation of the primary motor cortex depend on the amplitude of the motor evoked potentials. *PLOS ONE*, *12*(9), e0184910. https://doi.org/10.1371/journal.pone.0184910

13.	Petrichella, S., Johnson, N., & He, B. (2017). The influence of corticospinal activity on TMS-evoked activity and connectivity in healthy subjects: A TMS-EEG study. *PLOS ONE*, *12*(4), e0174879. https://doi.org/10.1371/journal.pone.0174879

14.	Oostenveld, R., Fries, P., Maris, E., & Schoffelen, J.-M. (2011). FieldTrip: Open Source Software for Advanced Analysis of MEG, EEG, and Invasive Electrophysiological Data. *Computational Intelligence and Neuroscience*, *2011*, 1–9. https://doi.org/10.1155/2011/156869

15.	Fischl, B. (2012). FreeSurfer. *NeuroImage*, *62*(2), 774–781. https://doi.org/10.1016/j.neuroimage.2012.01.021

16.	Van Veen, B. D., Van Drongelen, W., Yuchtman, M., & Suzuki, A. (1997). Localization of brain electrical activity via linearly constrained minimum variance spatial filtering. *IEEE Transactions on Biomedical Engineering*, *44*(9), 867–880. https://doi.org/10.1109/10.623056

17.	Westner, B. U., Dalal, S. S., Gramfort, A., Litvak, V., Mosher, J. C., Oostenveld, R., & Schoffelen, J.-M. (2022). A unified view on beamformers for M/EEG source reconstruction. *NeuroImage*, *246*, 118789. https://doi.org/10.1016/j.neuroimage.2021.118789

18.	Ermolova, M., Kozák, G., Belardinelli, P., & Ziemann, U. (2025). Plasticity of interhemispheric motor cortex connectivity induced by brain state-dependent cortico-cortical paired-associative stimulation. *Scientific Reports*, *15*(1), 32677. https://doi.org/10.1038/s41598-025-20294-1

19.	Glasser, M. F., Coalson, T. S., Robinson, E. C., Hacker, C. D., Harwell, J., Yacoub, E., Ugurbil, K., Andersson, J., Beckmann, C. F., Jenkinson, M., Smith, S. M., & Van Essen, D. C. (2016). A multi-modal parcellation of human cerebral cortex. *Nature*, *536*(7615), 171–178. https://doi.org/10.1038/nature18933
